# Creation of a High-Resolution, Continually Evolvable Lineage Tracer

**DOI:** 10.64898/2026.09.30.754670

**Authors:** Ryan N. Delgado, Yichen Si, Rebecca E. Andersen, ChangHee Lee, Henry L. Bushnell, Jordan L. Doman, Smriti Pandey, Alexander A. Sousa, Emma R. West, Carina E. Dagotto, David R. Liu, Constance L. Cepko

## Abstract

Cellular diversity is required for the proper function of many tissues, including the retina, with its ∼130 cell types.^1-3^ Previous lineage studies suggested that retinal progenitor cells (RPCs) generate cell diversity through biased cell divisions.^4-9^ However, the extent to which each progenitor cell contributes to diversity, and the overarching rules governing the lineages of the collection of progenitor cells, are unknown. While classic lineage tracing approaches including retroviral labeling^4,6,10^, dye injection^11,12^, and time-lapse microscopy^13,14^ were used to make the foundational insights regarding retinal lineages, they cannot be used to interrogate multigenerational lineages at scale. Here we describe the creation of an evolvable lineage tracing tool, “SCRIBBLE” for <u>S</u>equential <u>C</u>ombinatorial <u>R</u>ecorder for Iterative <u>B</u>arcode-<u>B</u>ased <u>L</u>ineage <u>E</u>volution. SCRIBBLE uses prime editor to iteratively edit a barcode sequence while preserving the order of genesis of clonally related progeny. Through a series of rationally designed high-throughput screens and design optimizations, we identified novel sequences that support a high level of barcode editing and information content. Evaluation of SCRIBBLE in embryonic mouse retina recapitulated known lineages in retinal development, demonstrating that SCRIBBLE is capable of high-resolution lineage tracing in a complex tissue.

---

The retina is a remarkably complex part of the central nervous system. It not only detects light, but it transforms the patterns of light into features that are useful to an organism, then transmits these features to the brain.^15^ In order to create these transformations, the retina relies on its immense cellular diversity. There are six principal classes of retinal cell types, but the total diversity of the retina is much more, with several classes comprised of many functionally distinct subtypes. Current estimates suggest there are ∼130 different subtypes of retinal cells in mice with most subtype diversity found in the interneurons, bipolar cells (15 subtypes)^2^ and amacrine cells (63 subtypes)^16^, as well as in the output neurons, the retinal ganglion cells (45 subtypes)^17^. The overarching strategy employed by the retina to produce such a diverse population of cells is largely unknown.

Studies of cell lineage – the series of cell divisions made by a progenitor cell and its progeny, and the cell types that derive from such divisions – have provided key insights into the developmental strategies used in the retina to generate cell diversity. These studies reveal a hierarchical relationship among retinal progenitor cells (RPCs). Early embryonic RPCs are highly proliferative and multipotent, producing large clones with variable cell-type composition^4,5^. Statistical analysis of these large, multigenerational clones fails to reveal obvious patterns of biased cell type production. However, nested within these large, variable lineages are terminally dividing RPCs biased towards producing certain retinal cell types^6-8^. Congruent with these observations, scRNA-seq analysis of the developing retina reveals several distinct populations of RPCs which have been referred to as primary RPCs and neurogenic RPCs (NPCs).^18,19^ Primary RPCs correspond to the more proliferative, multipotent population of RPCs, while NPCs refer to the terminally dividing population of cells. Both progenitor populations exist throughout retinal development. In the embryonic retina, scRNA-seq analysis revealed two distinct types of NPCs, identified by expression of either *Olig2* or *Galanin (Gal)*.^18,20,21^ Previous lineage tracing studies have demonstrated that embryonic *Olig2*+ NPCs are biased toward producing cone photoreceptors and horizontal cells^6^ while *Gal*+ NPCs are biased toward producing amacrine cells and RGCs^9^. Thus, together, the two subpopulations of early embryonic NPCs are biased to produce all four classes of embryonically produced retinal cell types. These studies, and others, including one based upon high resolution mapping of the spatial and temporal patterns of bipolar subtype genesis^22^, led to a hierarchical model of retinal development. The model suggests that highly multipotent RPCs divide multiple times to form large, multigenerational lineage trees, with fate biases emerging at the tips of terminal branches. The model also predicts that there is an obligate order of genesis of each cell type, with the same order displayed by each clone. A test of this model requires the ability to identify each cell type generated by each cell division, which has so far not been possible.

Classic lineage tracing studies employed a variety of methods that labeled RPCs with a heritable marker that was passed to progeny. These studies used vital dyes^11,12,23^, retroviral delivery of transgenes^4,6,10^, time-lapse microscopy^13,14^, recombinase-mediated activation of reporter genes^9,24,25^, and even chimeras^5^. More recently, the use of short DNA sequences referred to as “barcodes” have increased the throughput of lineage tracing studies by allowing the simultaneous labeling of many progenitors at a single time. These DNA barcodes are often encoded in the 3’UTR of expressed transgenes making it possible to capture these sequences along with cellular mRNA in scRNA-seq experiments. Such analyses allow for the simultaneous recovery of cell identity and clonal relationships. Although these techniques have greatly advanced our understanding of development, they are limited in their ability to resolve the order in which cell types are produced within a clone, as the labels are static. To address this problem several groups have created a new generation of lineage tracing techniques which use “evolvable” barcodes. These approaches rely on the mutation of a barcode sequence, causing it to “evolve” over the course of development. Since the barcodes are heritable, the mutations made in each generation are passed to the subsequent generation. Computational approaches are then used to compare the similarity/dissimilarity of evolved barcode sequences to infer the relatedness of cells within clones, similar to how evolutionary biologists use genomic data to assemble phylogenetic trees.

Over the last decade, several evolvable barcode methods have emerged, establishing that evolvable barcodes can work.^26-28^ However, these early studies also made it clear that several improvements were needed. An ideal evolvable lineage tracer would: 1) allow for theoretically infinite barcode evolution, 2) not rely on deletion as a means of barcode evolution, and 3) make the order of evolution apparent. Furthermore, given the power of comparative genomics, it would be greatly beneficial if the method could apply to a wide range of developing tissues and species. Here, we describe the creation of a new lineage tracing tool called SCRIBBLE (<u>S</u>equential <u>C</u>ombinatorial <u>R</u>ecorder for Iterative <u>B</u>arcode-<u>B</u>ased <u>L</u>ineage <u>E</u>volution) which meets these design criteria. Through a series of *in silico* and *in vitro* screens, we identified novel sequences with which to drive barcode evolution, optimizing SCRIBBLE’s edit rate and barcode sequence diversity. Benchmarking studies using scRNA-seq in the mouse retina confirmed that SCRIBBLE is sufficient to reconstruct multi-generational lineage trees. Importantly, it recapitulated clonal patterns in the retina that had been observed using other lineage tracing methods. SCRIBBLE can now be used to identify, at scale, how lineages contribute to diversity during development.

## RESULTS

In order to recreate multigenerational lineage trees, we employed prime editor to sequentially and iteratively add new sequences to the end of a transcribed barcode (**Figure 1A**). This barcode is comprised of two elements: 1) a 20bp “Static ID” which uniquely labels each barcode, allowing for the discrimination of multiple barcodes that could label a single clone, and 2) a single full-length protospacer to which new sequences are added. Collectively, we refer to this as a SCRIBBLE barcode. Additions to the SCRIBBLE barcode cause it to “evolve” in a unidirectional manner with each new addition (**Figure 1B)**. The directionality is based upon additions at a single full-length protospacer sequence located at the “growing” end of the barcode (**Figure 1A,B**), hereafter referred to as the “active site”. As a result, edits added to the SCRIBBLE barcode occur colinearly to the order they were made. To accomplish this editing scheme, we designed epegRNAs that target the active site and encode a complex prime edit which 1) deletes the 3’ PAM-proximal bp of the active site and 2) inserts 24bp of new sequence. As a result, the active site targeted in each prime edit is rendered inactive; it cannot be targeted again as it no longer has the requisite full-length sequence or adjacent PAM motif needed to function as a protospacer. The newly inserted 24bp sequence is comprised of a 4bp barcode fragment followed by a new full-length active site positioned at the distal end of the barcode. Thus, each SCRIBBLE edit adds one unit of information and establishes a new active site at the growing end of the barcode. This allows the editing process to repeat. When a progenitor cell divides and produces two daughter cells, each daughter cell will inherit the same edited barcode, but each will then continue to edit the barcode independently, causing the sequence to diverge (**Figure 1C**). By calculating both the shared root and divergence of SCRIBBLE barcodes among all cells of a clone, we can determine which cells diverged more recently from a common ancestor. The sequential, iterative nature of SCRIBBLE edits facilitates the reconstruction of multigenerational lineage trees.

**Figure 1.**
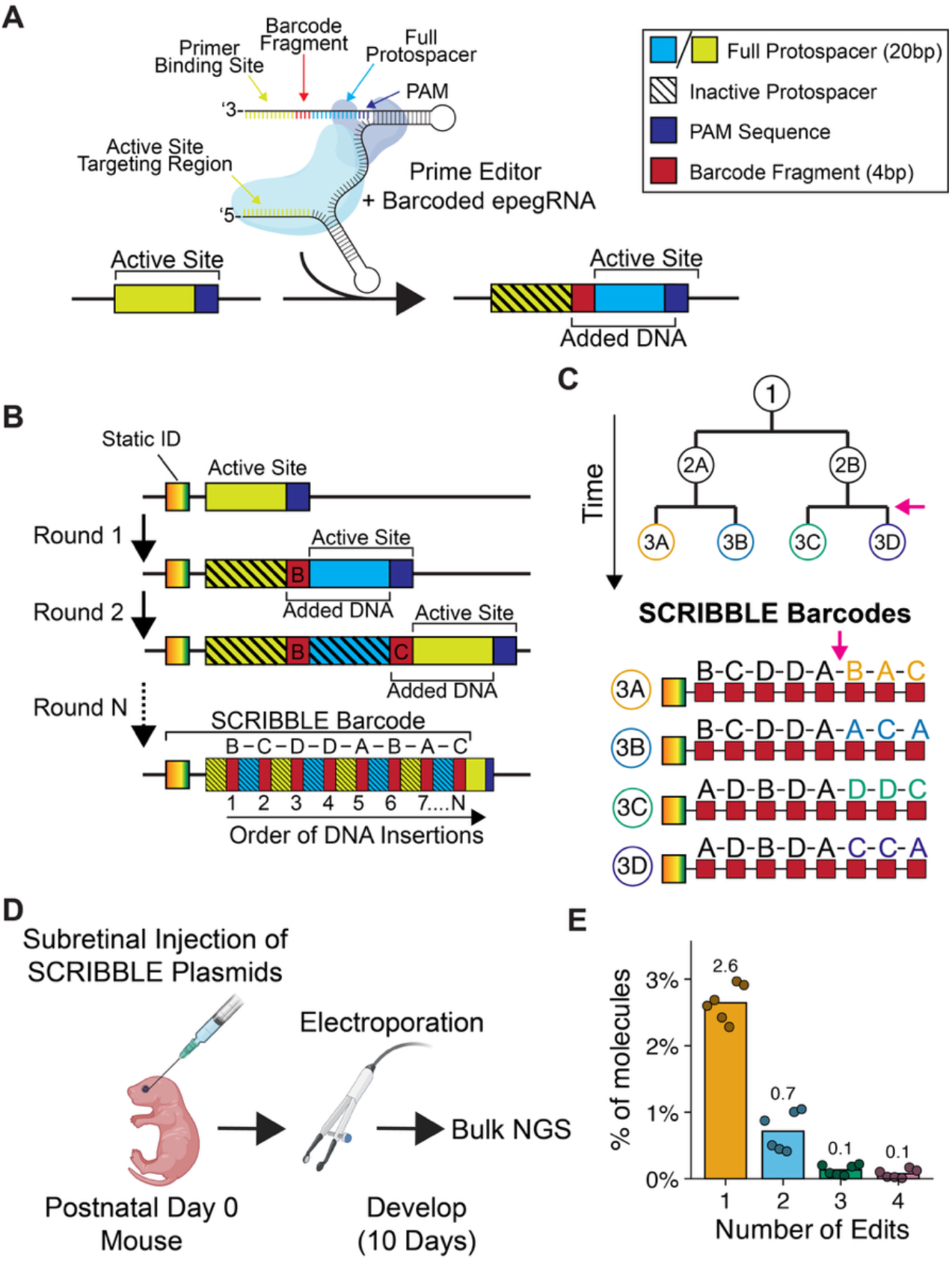
Conceptualization and Initial Testing of SCRIBBLE. **A)** Schematic depicting SCRIBBLE prime edit design including relevant elements of the SCRIBBLE barcode sequence. **B)** Schematic illustrating multiple rounds of SCRIBBLE editing and resulting evolutions to SCRIBBLE barcode sequence. **C)** Hypothetical three-generation lineage tree depicting series of cell divisions with cells depicted as circles. Below, corresponding hypothetical SCRIBBLE barcodes from the third generation of cells. Orange and purple arrows in lineage tree denote the second round of cell divisions at which time cells 3A and 3B diverge from each other and can acquire unique, distinguishing barcode edits after their shared common sequence. **D)** Experimental approach of SCRIBBLE prototype testing using Postnatal Day 0 mouse pups. **E)** Results of SCRIBBLE prototype testing depicted as a histogram showing the % of SCRIBBLE barcodes that were edited a given number of times, depicting only those with at least 1 edit. Individual dots depict results for biological replicates while bars depict mean.

To develop SCRIBBLE for *in vivo* lineage tracing, we designed a prototype which used a pair of active sites (*HEK3* and *VEGFA*) that had previously been successful targets of prime editor.^29^ They also had been shown to support DNA insertions (18bp) similar to the size of a SCRIBBLE edit (**Supplemental Figure 1A**). In addition to the active site, which is directly targeted by prime editor, the genomic sequence surrounding the active site can influence the efficacy of a prime edit. We observed that the HEK3 locus was particularly amenable to prime editing and that a minimal amount of sequence flanking the endogenous HEK3 site was sufficient to sustain high rates of prime editing. This was true even when the site was positioned outside of the endogenous HEK3 locus (**Supplemental Figure 1B-D**). Addition of a PE3-style^29^ nicking sgRNA which targeted part of the non-edited flanking sequence located 90bp downstream of the active site further increased editing (**Supplemental Figure 1D**). Using these sequences, we created a set of piggyBac transposase-compatible DNA plasmids encoding 1) target sequences containing an active site to initiate the evolution of SCRIBBLE barcodes, 2) epegRNAs that are used to make edits, and 3) a nicking sgRNA to increase the efficiency of editing. We delivered this pool of plasmids along with piggyBac transposase by electroporation to the postnatal day 0 (P0) mouse retina and allowed the retina to develop until P10 (**Figure 1D**). Analysis of barcodes by next generation sequencing (NGS) revealed iterative barcode edits, indicating that the prototype was functioning as intended (**Figure 1E**). However, only a relatively small number of all the recovered barcodes were edited (**3.6+/-0.7**%, mean ± S.D.) which is likely insufficient for high resolution lineage tracing. Thus, while certain features of the SCRIBBLE design, such as the flanking sequences used in the target sequence, had been methodically tested, other aspects including the active sites, barcode fragments, and nicking site remained as potential sources of improvement.

To improve SCRIBBLE edit rate, we performed a series of three screens (one *in silico*, two *in vitro*) aimed at identifying active sites that support high rates of prime editing. First, we analyzed all potential active sites by passing them through a series of filters (**Supplemental Figure 1E-G**, Methods) aimed at identifying novel sequences that are not present in the genomes of model organisms, thereby minimizing the chance that SCRIBBLE might edit an endogenous sequence, creating difficulty in clonal reconstruction and/or create mutations of endogenous genes. These model organisms included mouse, chicken, and human (**Supplemental Figure 1F**). After alignments and additional QC filters, there were ∼2.8 million sequences that could potentially serve as active sites. To further reduce the pool of putative sequences and prioritize the investigation of those likely to support high rates of prime editing, we calculated each sequence’s Doench 2016 score^30,31^, an on-target Cas9 efficiency score that we reasoned could be a surrogate for prime editing efficiency (**Supplemental Figure 1G**). We took the top scoring sequences (998) and validated them using an *in vitro* screen to determine which functioned well as an active site that could be targeted by prime editor. To accomplish this, we designed a simple prime editing scheme in which we targeted each sequence with the same 18bp insertion using prime editor (**Figure 2A**). Using a similar barcode design as in the pilot including the HEK3 locus flanking sequences, we replaced the HEK3 active site with our top scoring sequences, cloned in the matching epegRNA to perform the prime edit, and delivered the resulting pool of molecules in lentiviral format to HEK293T cells (**Figure 2A**). NGS analysis revealed that 220 sequences were edited to greater levels than the HEK3 protospacer itself indicating it is possible to improve upon prime editing efficiency from an already highly permissive genomic context (**Figure 2A**). On average, the top 12 rank-ordered sequences scored **68%** greater compared to the HEK3 control and were chosen for further development. We also identified three high-scoring sequences as potential replacements for the previously described endogenous HEK3 nicking guide used to facilitate PE3-style edits in the SCRIBBLE barcode (Methods).

**Figure 2.**
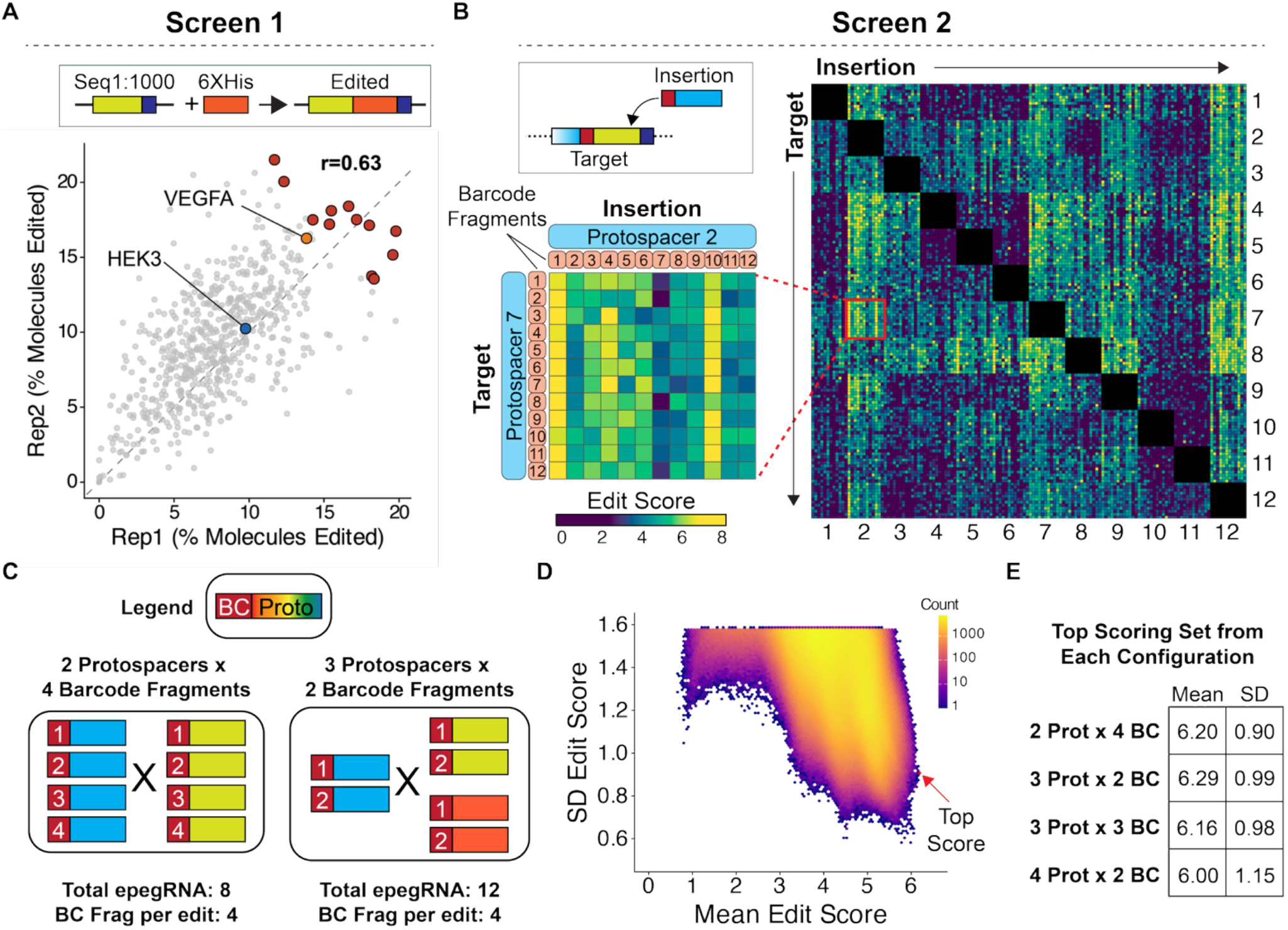
Rationally Designed, High-Throughput Screens Identify Optimized SCRIBBLE Sequences. **A)** Top, Prime editing scheme used in Screen 1 in which a 18bp 6xHis tag was inserted into each of 1000 protospacer sequences. Bottom, Results of Screen 1 depicted as a scatter plot showing the percentage of edited molecules in two experimental replicates (Replicate 1 on X-axis, Replicate 2 on Y-axis). Individual protospacers depicted as dots. Red dots, top 12 most highly scoring protospacers. Orange dot, endogenous human VEGFA protospacer. Blue dot, endogenous human HEK3 protospacer. **B)** Top left, Prime editing scheme used in Screen 2 in which top 12 scoring protospacers from **Figure 2A** were inserted into active sites and also served as active sites. Right, Results of Screen 2 depicted as a heatmap showing Edit Scores (see methods) for every pairwise combination of tested sequences as both targets (rows) and insertions (columns). Heat plot is organized first by protospacer sequences in numeric order (1:12) with barcode fragments (1:12) nested within each protospacer. Bottom left, Inset of Screen 2 results depicting high edit scores between Protospacer 7 containing target sites and Protospacer 2 containing insertions. **C)** Schematic depicting two potential SCRIBBLE configurations which use either two or three protospacers for iterative, sequential barcode editing. Different barcode fragments shown as a number in red box while different protospacers depicted as solid color. Total number of epegRNAs required for each configuration and number of potential barcode fragments added per edit listed below. **D)** Density plot showing mean and standard deviation Edit Score for every possible combination of measured edits from Screen 2 that fit a 2 Protospacer x 4 barcode fragment configuration. Top scoring combination depicted with red dot and arrow. **E)** Mean and SD scores from the top scoring sets of different protospacer/barcode fragment configurations.

While this first screen identifies potential active sites on the basis of their ability to be the target of a prime edit, SCRIBBLE barcode evolution requires that inserted sequences themselves are sites of efficient insertion by prime editor. Previous studies have demonstrated that prime editing efficiency can vary widely depending on the edit, even when the edit is made to the same locus. Therefore, we reasoned that the active sites that best supported SCRIBBLE-style editing would not only have high individual prime editing efficiencies, but could also be paired to create the highest efficiency for sequential SCRIBBLE-style edits (**Figure 2B**). Therefore, we designed a second *in vitro* screen to calculate a pairwise edit score between every combination of potential active sites identified in the first screen. To more closely match an actual SCRIBBLE edit, we created a set of twelve 4bp barcode fragments and included them in the epegRNA design such that every edit would insert a barcode fragment-active site-PAM combination in exactly the way a SCRIBBLE edit would. We performed this screen in HEK293T cells and analyzed the resulting edited sequences by NGS, calculating a relative enrichment score for each potential edit (**Figure 2B**, Methods). Between biological replicates, we observed a correlation score of r=0.63, indicating that despite the complex nature of the screen, the results were reasonably reproducible. SCRIBBLE barcode evolution requires at least two active sites and a pool of potential barcode fragments in order to create diverse barcode sequences. While increasing the number of barcode sequences that can be inserted at each round of editing would improve the resolution of lineage reconstruction, it also increases the number of epegRNAs needed (**Figure 2C**). Creation and delivery of a highly diverse barcoded epegRNA set can be technically challenging. We therefore sought to identify several high-efficiency “configurations” of active sites and barcode fragments that provided sequence diversity with a reasonable number of epegRNAs. Using the results of our pairwise screen, we calculated the mean edit score for every possible combination of sequences that matched four different configurations. These results demonstrated that top-scoring sets in different configurations exhibited similar edit scores, suggesting that any of these top-scoring sets might support high editing rates (**Figure 2D, E)**.

To validate these screen results, we tested the top scoring sets in mouse retinal explants (**Figure 3A**). First, we tested the top scoring 2- and 3-active site configurations in combination with the three novel PE3-style nicking sites identified in the first screen (**Figure 3A**). All combinations demonstrated iterative SCRIBBLE barcode editing and those containing a nicking site had ∼2-fold more edits than combinations that did not have PE3-style editing (**Supplemental Figure 2A,B**). Nicking site 3 (referred to as “Nick3”) slightly outperformed the other nicking sites across all conditions with **15±7.2%** (mean ± S.D., n=4) sites edited in the 2-protospacer configuration and **25±7.1%** (mean ± S.D., n=4) sites edited in the 3-protospacer configuration. The difference in overall edit rate between these two configurations came mostly from targets that had been edited a single time (**9.1±3.7%** in 2-protospacer vs. **18.7±4.3%** in 3-protospacer, **Supplemental Figure 2A**,**B**). While a direct comparison to the initial SCRIBBLE prototype prior to these optimizations (**Figure 1E**) is not possible given the inherent differences in how the assays were performed (10-day *in vivo* vs 5-day explant tracing), these results demonstrate the novel SCRIBBLE sequences support iterative editing and a promising edit rate. We decided to retain Nick3 in the SCRIBBLE design for all future experiments.

**Figure 3.**
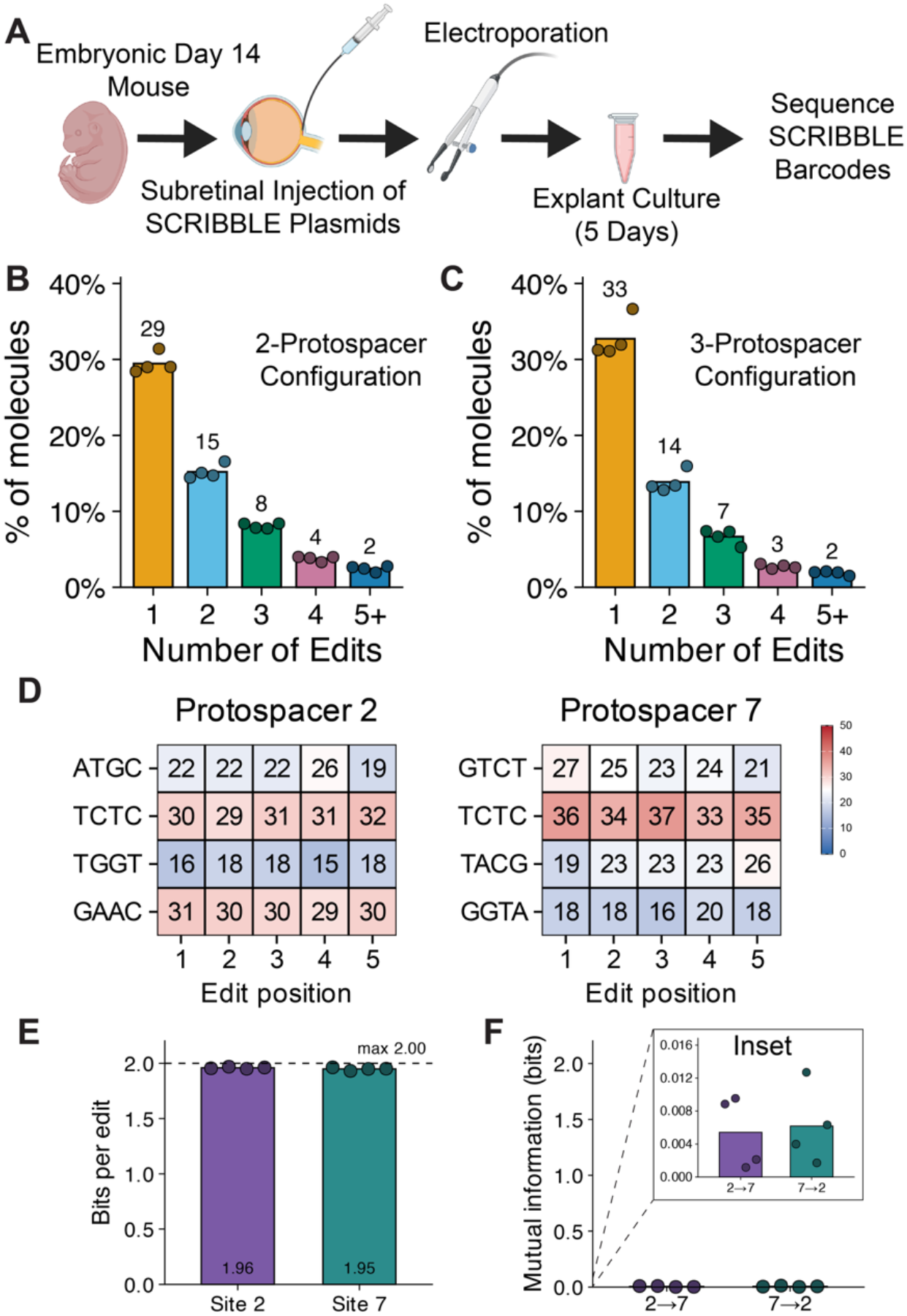
Test of Optimized SCRIBBLE Sequences in Mouse Retinal Explants. **A)** Schematic depicting experimental approach for the testing of optimized SCRIBBLE sequences in Embryonic Day 14 mouse retinas. **B-C)** Results of **(B)** two-protospacer or **(C)** three-protospacer optimized SCRIBBLE configuration depicted as a histogram showing the percentage of SCRIBBLE barcodes edited a given number of times, depicting only those with at least 1 edit. Individual dots depict results for biological replicates while bars depict mean. **D)** Heatplot showing relative abundance of each barcode fragment across edit positions 1 through 5 in from two-protospacer configuration shown in **Figure 3B**. Barcode fragments depicted across rows and edit positions across columns. Values are normalized to edit position. **E-F)** Shannon entropy **(E)** and Mutual Information **(F)** calculated for all edits made to Protospacer 2 or Protospacer 7 target sites. Mean values depicted as a bar while biological replicated depicted as dots. Dashed line in (**E**) depicts the theoretical maximum entropy of a 4 barcode fragment design. Inset in (**F**) included in order to better visualize data which is compressed on y-axis when scaled to 2.0 bits.

Next, we tested whether several features that were found to generally improve prime editing could further enhance SCRIBBLE editing. A previous study of prime editor optimization found that use of the Flip + Extension (F+E) scaffold^32^ sequence in epegRNA improved prime edit rates compared to the canonical scaffold sequence.^33^ All experiments described above had been performed using a modified canonical scaffold (referred to as sgRNA scaffold 2 in Anzalone et al.^29^) in which the second to last bp in the terminal hairpin are inverted. Exchanging this scaffold with the F+E scaffold resulted in a modest increase of ∼**1.3-fold** in edited SCRIBBLE barcodes for both the 2- and 3-active site configurations (**Supplemental Figure 2C,D**). We also tested JLD1907, a previously unreported prime editor variant from the phage-assisted evolution campaign that produced the suite of PE6 editors.^34^ JLD1907 was characterized during that campaign but was not included in the named PE6a-g suite, as it did not improve editing broadly across edit types and target sites; instead it performed well in a subset of sequence and edit contexts, consistent with the specialization of evolved prime editors reported in that study.^34^ Reasoning that the fixed insertion and small set of defined active sites in SCRIBBLE constitute such a context, we tested JLD1907 in mouse retinal explants and observed a 2.5-fold increase in editing compared to PEmax (**Supplemental Figure 2E**).^34^To determine if this optimized SCRIBBLE and JLD1907 would also work in other species, it was tested in the chick retina from E2 through E10. This test revealed substantial SCRIBBLE barcode evolution, demonstrating that the optimized sequences work in different species (**Supplemental Figure 2F**). Thus, SCRIBBLE editing can be further improved by implementing more general improvements to prime editing.

Single cell RNA sequencing in cells with SCRIBBLE barcode evolution would allow for the simultaneous recovery of cellular mRNA and SCRIBBLE barcodes so that both cellular identity and lineage relationships could be determined. Our experiments described above used analysis of SCRIBBLE barcode editing at the DNA level (Methods). To test whether SCRIBBLE barcodes could be transcribed into RNA and then recovered as they would in an actual lineage tracing experiment, we repeated the experiment using a 2-protospacer configuration, JLD1907, and sgRNA scaffold 2 (as described in **Supplemental Figure 2E**), followed by sequencing of cDNA (Methods). Recovery of RNA was possible as the integrated SCRIBBLE target constructs included a promoter sequence. We were able to successfully recover transcribed SCRIBBLE barcodes. This analysis revealed a **1.6-fold** increase in the amount of edited SCRIBBLE barcodes, compared to those recovered from genomic DNA (**Figure 3B**). We observed similar levels of SCRIBBLE barcode edits in the 3-protospacer configuration using cDNA (**Figure 3C**). The observed differences in SCRIBBLE edits at the genomic DNA and cDNA (RNA) level in the same experimental condition might be explained by previous studies which found that more euchromatic, and therefore more highly transcribed, regions of the genome support higher levels of prime editing.^35^ This result would suggest that the most edited SCRIBBLE barcodes would originate from more highly expressed loci, and therefore would be more likely to be captured by scRNA-seq, in addition to being more efficiently edited.

Qualitative analysis of the results from the optimized SCRIBBLE revealed diverse sequence evolution. In its current form, each SCRIBBLE edit has the possibility of incorporating one of four different barcode fragments. Analysis of evolved barcode sequences from **Figure 3B** (2-protospacer configuration, sgRNA scaffold 2, RNA) revealed that addition of each barcode fragment occurred frequently and at near even frequency (**Figure 3D**). The information content (Shannon entropy) of SCRIBBLE was ∼1.95 bits for each edit which is close to the configuration’s theoretical maximum of 2.0 bits (**Figure 3E**). Furthermore, we found that sequential barcode fragment addition occurred randomly such that the preceding barcode fragments did not bias subsequent edits (**Figure 3F** and **Supplemental Figure 2H**). Together the improvements made to SCRIBBLE now support high rates of editing, resulting in highly diverse, evolved sequences suitable for reconstructing multigeneration lineage trees.

To test the optimized SCRIBBLE tool in the developing embryonic mouse retina, we performed lineage tracing in early retinal development starting at embryonic day 12.5 (E12.5) (**Figure 4A**). At this time, retinal ganglion cells, amacrine cells, horizontal cells, and cone photoreceptors are being produced.^36^ Electroporation was used to deliver SCRIBBLE components to E12.5 explants, and explants were cultured for 3 days. As GFP was included on the PE plasmid, GFP+ cells were isolated by On-Chip Sort microfluidics device and used for scRNA-seq. Three different biological replicates were pooled for this analysis (Methods). During preparation of the transcriptomic library, we added a spike-in primer to sub-amplify captured SCRIBBLE barcode reads to generate a SCRIBBLE sequencing library (Methods). Analysis of transcriptomic data revealed the four principal classes of post-mitotic retinal neurons born during this time, as well as several populations of RPCs (**Figure 4B,C**). Barcodes were recovered from 68% of cells, as expected from the GFP enrichment (**Supplemental Figure 3A**, Methods). Recovery of SCRIBBLE barcodes was even across different cell types suggesting that there was unlikely to be cell-type specific biases in recovery of SCRIBBLE barcodes (**Supplemental Figure 3B)**. From these data, we were able to identify 852 multi-cellular clones which were defined by having at least 2 unique Static ID sequences (**Figure 4D**). Since individual clones can be labeled with multiple insertions of SCRIBBLE barcodes, the collection of all Static ID sequences within a clone serves as a “Clone ID”. Multi-cellular clones ranged in size from 2 to 5 cells with the majority (82%) of clones containing 2 cells (**Figure 4E**). These clone sizes are congruent with expected sizes considering the delay in barcode integration. This delay is due to the requirement for transcription and translation of piggyBac tansposase for integration of SCRIBBLE components, as well as cell loss from sample preparation for scRNA-seq.

**Figure 4.**
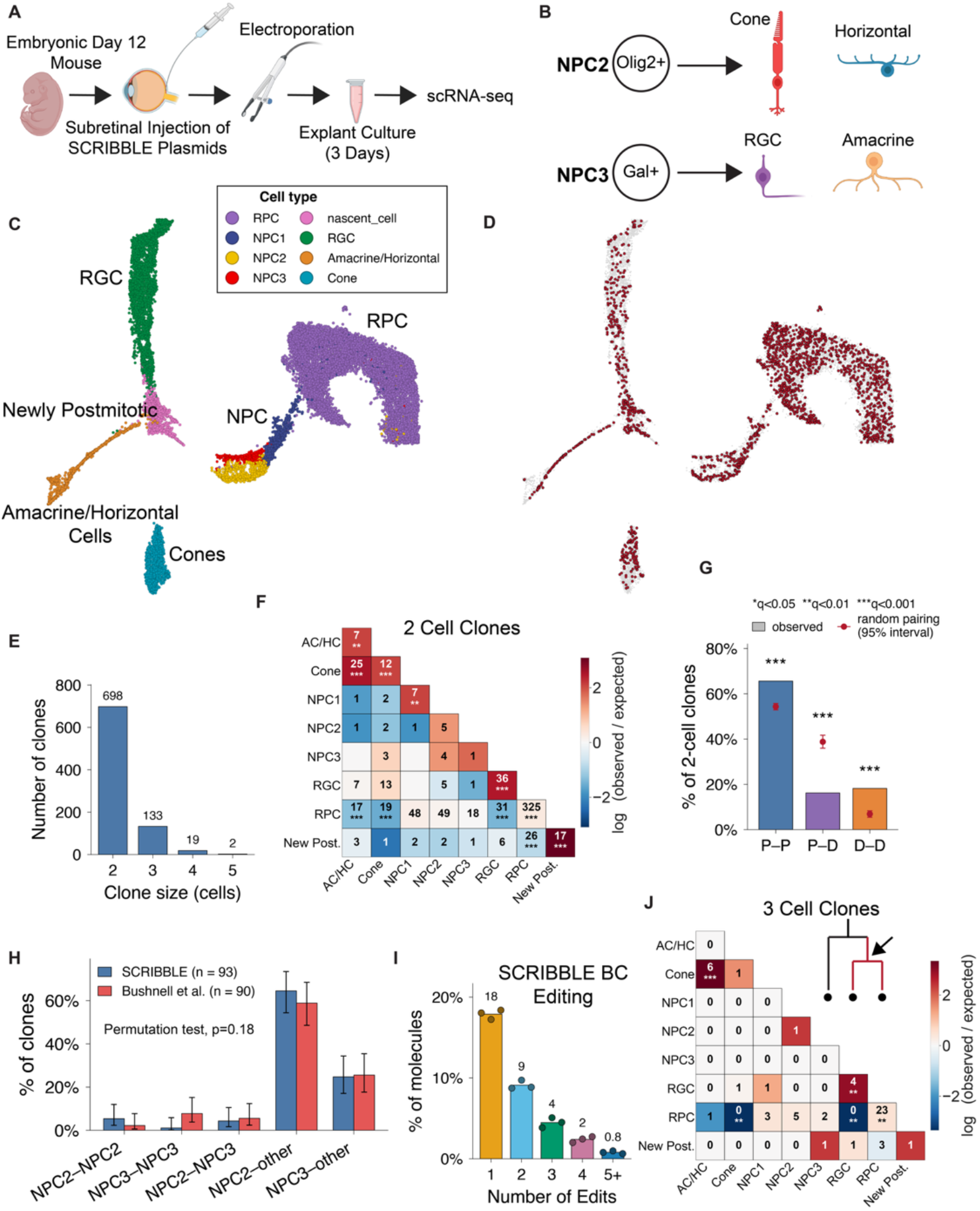
Optimized SCRIBBLE Recapitulates Known Lineages in Embryonic Retina. **A)** Experimental approach used to test optimized SCRIBBLE using E12.5 mouse retinas. **B)** Schematic depicting known biases in terminally dividing neurogenic progenitor cells (NPCs) in embryonic mouse retina. Each division can produce both heterotypic or homotypic combinations of depicted cells. **C)** UMAP plot depicting major retinal cell classes from captured cells. **D)** UMAP plot depicted cells that are part of multi-cellular clone defined by SCRIBBLE barcode highlighted in red. **E)** Histogram depicting clone size of reconstructed clones. **F)** Heatmap showing all pairwise co-occurrences of major cell classes in two-cell clones. Number in each cell depicts the number of clones in which a combination was observed and the color of each cell depicted the log(observed/expected) value. **G)** Percentage of two-cell clones that contained proliferative cells (P, i.e. progenitors) or differentiated cells (D, i.e. post-mitotic cells). Observed frequency of P-P, P-D, and D-D combinations shown as bars while mean and 95% range of expected frequencies as estimated by Monte Carlo simulation shown as red dot and bars. **H**) Bar plot comparing the percentage of two-cell clones containing at least one NPC2 or NPC3 cell in SCRIBBLE data (blue bars) to Bushnell et al, 2026 dataset derived using orthogonal Cre-based lineage tracing approach. Error bars depict a 95% Wilson confidence interval. The two distributions did not differ significantly (Monte Carlo permutation test, 200,000 permutations, p = 0.18). **I)** SCRIBBLE editing in multi-cellular clones depicted as a histogram showing the percentage of SCRIBBLE barcodes that were edited a given number of times, depicting only those with at least 1 edit. Individual dots depict results for biological replicates while bars depict mean. **J)** Heatmap showing all pairwise co-occurrences of major cell classes of inner branch of three-cell clones. Number in each cell depicts the number of clones in which a combination was observed and the color of each cell depicted the log(observed/expected) value. (F, J) Stars denote Benjamini– Hochberg-corrected significance from 100,000 label-shuffling permutations (two-sided): *q < 0.05, **q < 0.01, ***q < 0.001.

To determine if the cellular composition of SCRIBBLE clones was consistent with prior observations of cell type production from RPCs and NPCs from this age, two cell clones were compared to data from previous lineage studies. We observed that RGCs, cone photoreceptors, and amacrine/horizotonal cells were statistically more likely than expected to occur in homotypic clones (**Figure 4F**), consistent with previous lineage tracing studies^6,9^. More generally, we found these clones to result from either proliferative or differentiative divisions in which both members of the clone were either RPCs or post-mitotic cells (**Figure 4G**). Mixed clones containing one RPC and one postmitotic cell were significantly less common than expected by chance (**Figure 4F,G**). These results are consistent with the rapidly growing state of the early retina driven by the output of both proliferative (i.e. RPC) and terminally dividing progenitors (i.e. NPCs). Additionally, the lineage relationship of *Olig2*+ NPCs (NPC2) and *Gal*+ NPCs (NPC3), which were recently reported using a cre-based in vivo approach, were indistinguishable from SCRIBBLE-labeled clones (**Figure 4H**). The majority of these two NPC subtypes were previously found to occur in 2 cell clones with a non-NPC sibling (**Figure 4H)** ^9^ and the results from SCRIBBLE indicate that this sibling is frequently a RPC (**Figure 4F**). Collectively, these results demonstrate that SCRIBBLE faithfully recapitulates known biases in retinal cell development.

An analysis also was conducted to ask whether SCRIBBLE could reconstruct multi-generational lineage trees in the retina, i.e. clones > 2 cells. Analysis of SCRIBBLE barcodes within multi-cellular clones demonstrated a wide range in barcode number with 17.6% (150/852) having 10 or more SCRIBBLE barcodes (**Supplemental Figure 3D**). SCRIBBLE barcodes that defined multi-cellular clones were edited multiple times (**Figure 4I**) to an extent that was consistent with that observed in the bulk retinal experiment (**Figure 3B**), which had developed for an additional 2 days. Complete reconstruction of all cells within a clone was successful for 41% (54/133) of three cell clones and 21% (4/19) of four cell clones. Although the number of clones with three or more cells is limited in this benchmarking study, it was possible to determine if known biases in retinal development were represented in larger clones. Focusing specifically on the cells located on the inner branch of three-cell clones, the same enrichment analysis as was done with two-cell clones was carried out (**Figure 4J**). Here, the same observation of a bias for RGCs and RPCs to occur in homotypic pairs was made. Additionally, cone photoreceptors were found to occur together with cells in the amacrine/horizontal cell cluster more than would be expected by chance, consistent with what was observed in two-cell clones (**Figure 4F**). While we cannot transcriptionally distinguish between amacrine cells and horizontal cells at this developmental stage, it is likely that the cell type in these clones were horizontal cells, as previous lineage tracing of the progeny of Olig2+ NPCs demonstrated their bias to produce heterotypic pairs of cones and horizontal cells, along with homotypic pairs of cones. Also consistent with biases observed in our analyses of two-cell clones, RPCs within the inner branch were biased against being paired with post-mitotic cells, such as RGCs or cone photoreceptors, and biased toward being paired with other RPCs (**Figure 4J**). Thus, these data recapitulate the patterns seen in previous lineage studies of retinal development that used orthogonal methods. SCRIBBLE can thus be used to successfully recreate multi-generation lineage trees.

## DISCUSSION

In this study, we describe the design and optimization of SCRIBBLE, an evolvable lineage tracing tool. Through an iterative series of design improvements and high-throughput screens (**Figure 2**), we successfully identified novel protospacer sequences and barcode fragments that could facilitate high rates of iterative prime editing, leading to the evolution of diverse barcode sequences (**Figure 3**). Testing of the optimized SCRIBBLE design in primary retinal tissue recapitulated known biases of RPCs, including those of terminally dividing NPCs and more proliferative RPCs (**Figure 4**). While the duration of lineage tracing in this experiment was brief, it was sufficient to validate the utility of SCRIBBLE. The wide distribution of barcode number defining each clone suggests that piggyBac transposition of SCRIBBLE barcodes and other SCRIBBLE components is likely over-dispersed with some clones receiving many transpositions and others receiving few (**Supplemental Figure 3D**). Future improvements to SCRIBBLE would include approaches that increased the number of transpositions per clone or those that prospectively enriched for clones with a high transposition rate prior to capturing for scRNA-seq. Future studies using SCRIBBLE to mark thousands of clones generated over a longer period of time will be helpful in identifying the overarching strategy employed by the retina to generate its great diversity of cell types. In particular, it will allow a test of a model that we recently developed of a hierarchical clonal pattern, with an obligate order of genesis of retinal cell types in each clone.

Since we started developing SCRIBBLE, other groups have also developed evolvable lineage tracing tools. More recent “second generation” evolvable barcode tools have improved upon the common shortcomings of “first generation” evolvable barcode tools. One major area of improvement has been in how barcodes evolve over sequential rounds of editing. First generation tools drove barcode evolution through a variety of means that resulted in sequence evolution randomly throughout the barcode sequence, making it difficult to know the order in which the edits occurred.^26-28^ SCRIBBLE and other tools including DNA Typewriter^37^ and peCHYRON^38^ were designed so that edits are recorded sequentially within the growing barcode in the order in which they occur, helping reduce this ambiguity and increase the accuracy of lineage reconstruction. All three of these tools use prime editor. SCRIBBLE and peCHYRON differ from DNA Typewriter in that their design is open-ended allowing for theoretically infinite editing. In contrast, DNA typewriter is limited to a finite number of edits by design because it uses a “pre-patterned” barcode sequence composed of tandem repeats of a partial protospacer. A recent lineage tracing study of mouse embryogenesis using DNA typewriter with a capacity for six edits shows that while some barcodes are saturated after 13.5 days of editing, others still have the capacity to be edited again.^39^ Barcodes that saturate quickly are by definition highly amenable to being edited repeatedly and therefore likely to continue being edited, providing the highest resolution in reconstructing lineages. After five days of tracing in the mouse retina, ∼2% of SCRIBBLE barcodes have at least 5 edits (**Figure 3B,C**). Since SCRIBBLE is open-ended by design, these high performing barcode sites can continue being edited and further contribute to lineage reconstruction. While peCHYRON also uses an open-ended design, it differs from SCRIBBLE in its edit rate. Although we have not conducted a direct comparison of the two methodologies, and prime edit rates can differ between cell types, the current version of peCHYRON reports ∼2% of barcodes with 5 or more edits after 27 days of recording^38^ while SCRIBBLE has ∼2% of barcodes with 5 or more edits after 5 days of recording (**Figure 3B,C**). Given the magnitude of this difference, these two tools could be used to answer different biological questions— SCRIBBLE being used for rapidly dividing cell types in which high-resolution lineage reconstructions are important and peCHYRON for protracted events that take multiple months.

Other recent evolvable barcode tools have sought to increase the resolution of lineage reconstruction by increasing barcode density. For instance, PEtracer is a prime-editor based evolvable lineage tracing tool whose target sites can be edited a single time resulting in one of eight different states.^40^ By encoding 105 editable sites in a mouse ES line, they achieved a sufficient number of barcodes sites to perform lineage tracing for the first 10 days of mouse embryogenesis.^41^ Another tool called BASELINE uses a cas12a adenine base editor (ABE) to edit a large, 1.5kb barcode containing 50 target sites, converting adenine-thymine base pairs into guanine-cytosine base pairs.^42^ The technology leverages cas12a’s ability to process multiplexed guide RNAs which are used to target the 50 different target sites, fitting both guides and targets in a single plasmid. Hypercascade is another ABE-based lineage tracing tool that utilizes a large number of target sites, but achieves this in a more compact manner by utilizing a multi-layered design in which edits can generate deeper layers of target sites which can subsequently be edited. While the edits produced by ABE-based lineage tracers occur in a random order, are finite in number, and are informative only when there is diversity across cells in a clone, these tools utilize a large total number of target sites to off-set these inherent limitations.

The growing set of evolvable lineage tracing tools now offers experimentalists a diverse set of options to use. Depending on the biological question being studied, some tools might be better suited than others. So far, most of these tools have been described in proof-of-principle studies. As these tools start to be used to explore novel biology, the field as a whole will start to get a better sense of the role that lineage plays in disease and development.

**Supplemental Figure 1.**
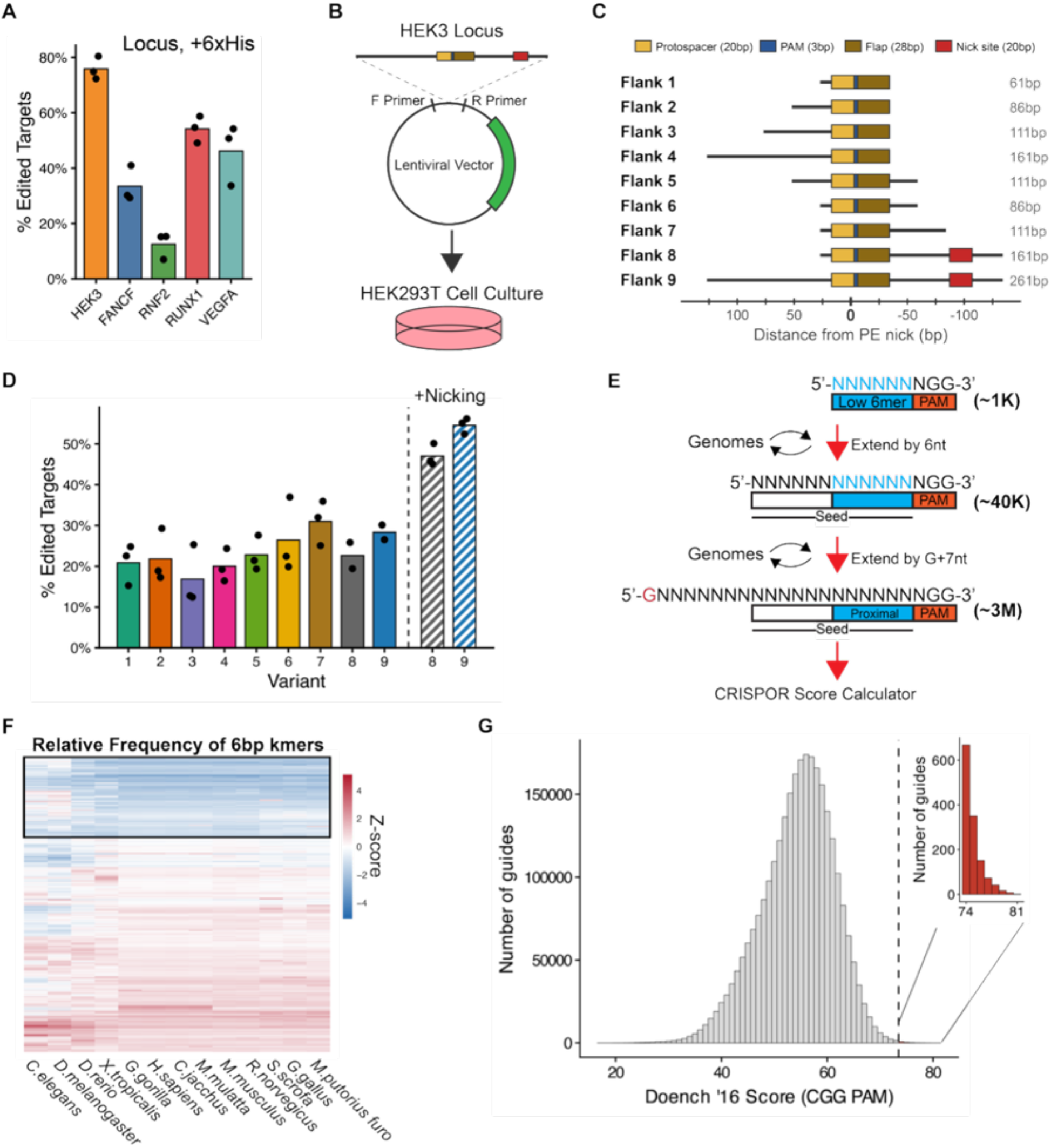
**A)** Bar plot depicting the percentage of targets with different protospacers edited with +6His insertion. Bar depicts mean and individual dots depict replicates. **B)** Schematic depicting design of lentiviral construct used to deliver variants of endogenous HEK3 locus sequence *in trans* to HEK293t cells. **C)** Description of HEK3 sequence variants tested and the amount of flanking sequence included. **D)** Results of HEK3 variant prime editing test depicting the percentage of molecules edited. Bar depicts mean and individual dots depict replicates. **E)** Iterative strategy used to create novel 20bp protospacer sequences not found in wide variety of genomes. The bold number to the right indicates the approximate number of sequences meeting each step’s criteria. **F)** Heat plot showing kmer frequency of all possible 6bp sequences across genomes of 13 model organisms. 6bp kmers with low z-scores used in subsequent iteration bound in black box. **G)** Results of *in silico* screen depicted as a histogram showing the Doench 2016 Scores of the ∼3M 20mers that passed all steps shown in panel E. Inset shows highest scoring sequences used for subsequent *in vitro* screen.

**Supplemental Figure 2.**
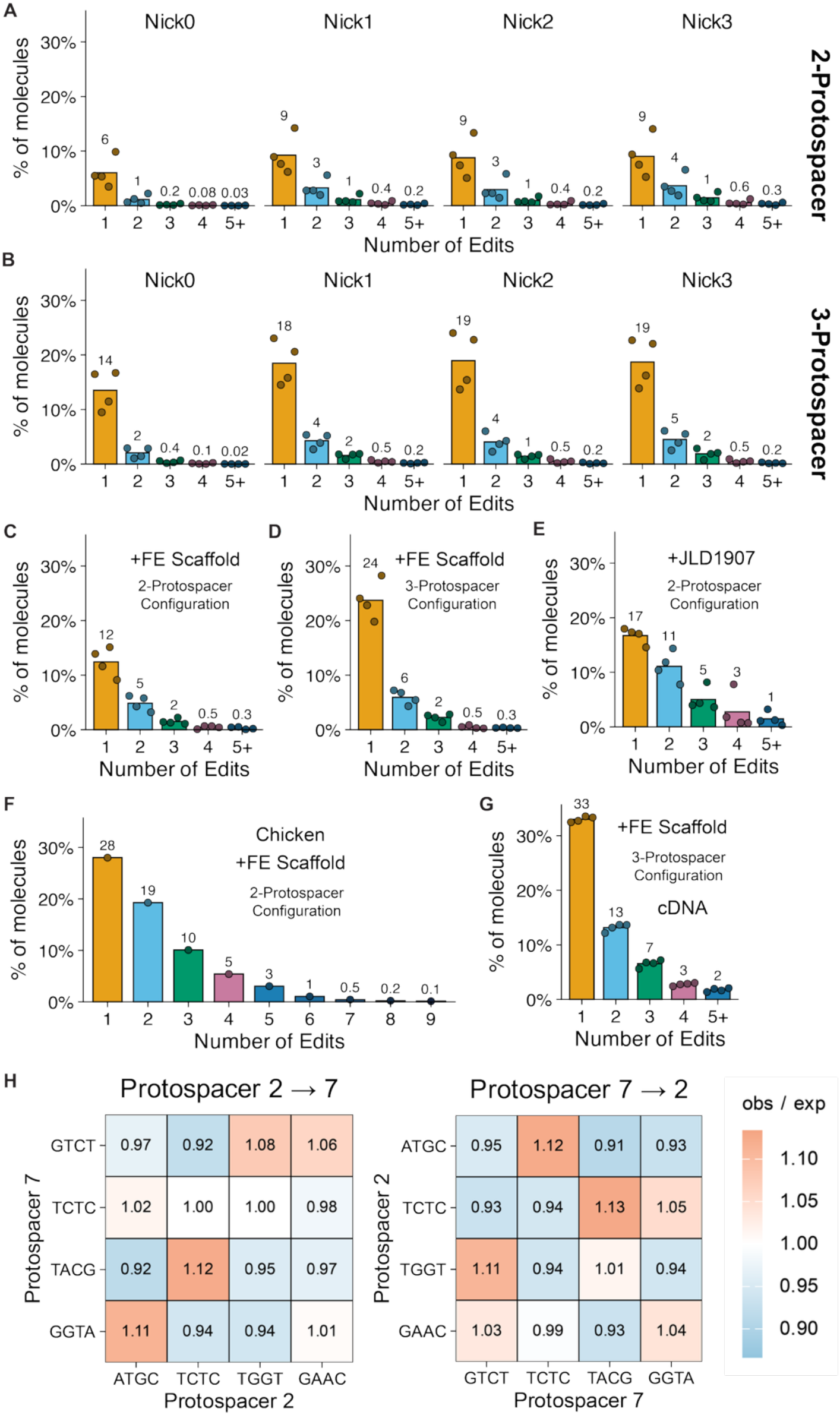
**A-G)** Results of various configurations of optimized SCRIBBLE testing depicted as a histogram showing the % of SCRIBBLE barcodes that were edited a given number of times, depicting only those with at least 1 edit. Individual dots depict results for biological replicates while bars depict mean. **(A)** Two-protospacer, sgRNA Scaffold 2, PEmax, DNA. **(B)** Three-protospacer, sgRNA Scaffold 2, PEmax, DNA. **(C)** Two-protospacer, F+E Scaffold, PEmax, DNA, **(D)** Three-protospacer, F+E Scaffold, PEmax, DNA, **(E)** Two-protospacer, sgRNA Scaffold 2, JLD1907, DNA, **(F)** E2 to E10 chicken retina, Two-protospacer, F+E Scaffold, JLD1907, DNA, **(G)** Three-protospacer, F+E Scaffold, PEmax, RNA. H) Heat plot depicting observed/expected ratio of sequential SCRIBBLE barcode edits. The rows in each panel correspond to different barcode fragments being targeted and the columns correspond to barcode fragments being inserted.

**Supplemental Figure 3.**
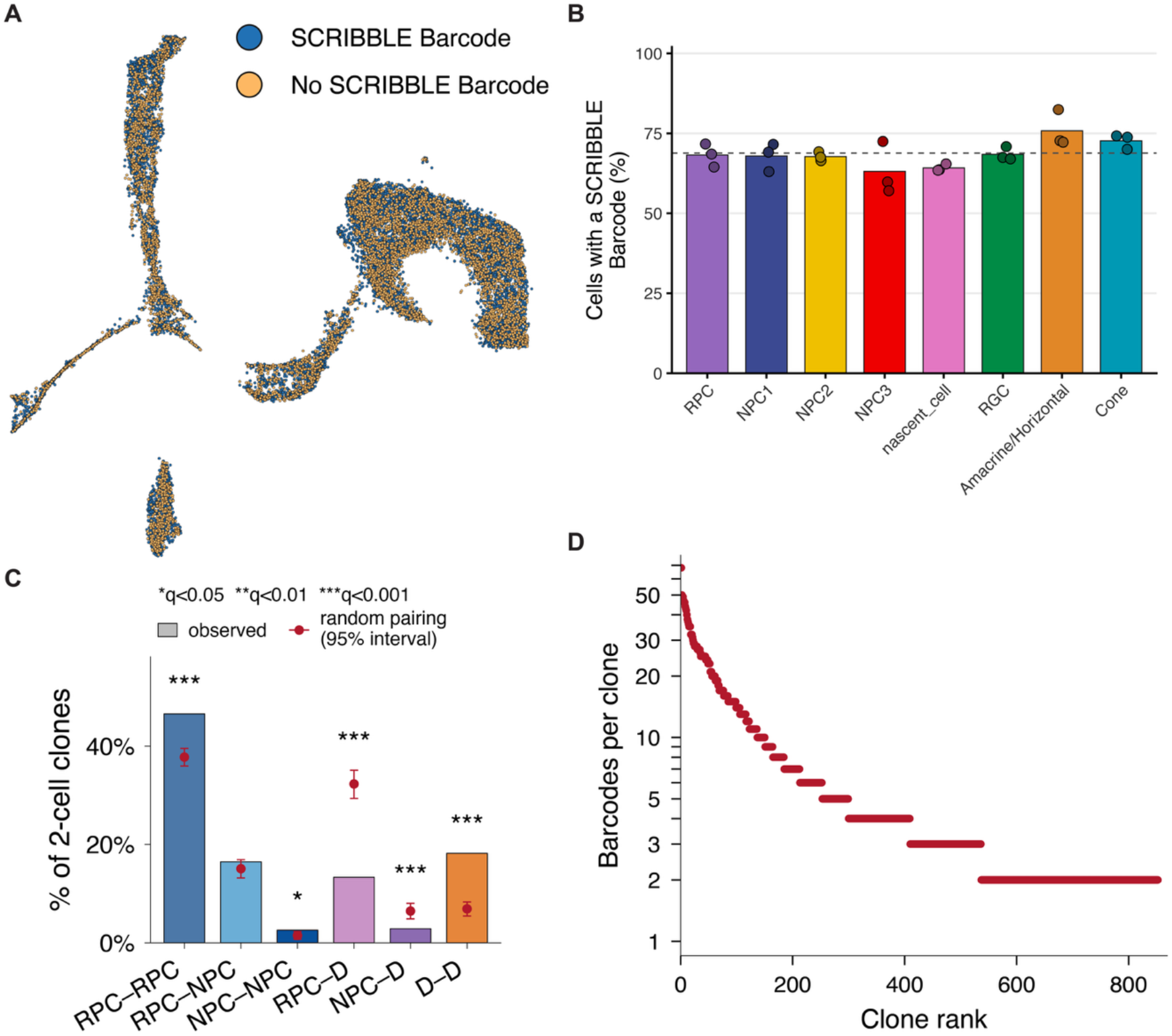
**A)** UMAP plot depicted cells that contain a SCRIBBLE barcode in blue and those without a SCRIBBLE barcode in orange. **B)** Bar plot depicting the percentage of retinal cell classes that have a SCRIBBLE barcode. Individual values from each biological specimen depicted as dots and un-weighted mean values across all three samples as bars. Dashed horizontal line depicts the overall average number of cells with a SCRIBBLE barcodes across the entire dataset. **C)** Percentage of two-cell clones that contain RPCs, NPCs, or differentiated cells (D, i.e. post-mitotic cells). Observed frequency of all pairwise combinations of these classifications shown as bars while mean and 95% range of expected frequencies as estimated by Monte Carlo simulation shown as red dot and bars. **D)** Rank order plot depicting the number of unique Static IDs used to define each clone (red dot). Y-xis is log10-scaled.

## MATERIALS AND METHODS

### Molecular Cloning

Most molecular constructs were generated by standard cloning techniques including restriction enzyme and ligation-based cloning as well as Gibson reaction.1 Constructs were transformed in chemically competent e. coli (DH5α prepared using Zymo, #T3001) and amplified by culture in a bacterial shaker by growing overnight at 37C. Constructs with high amounts of repeated elements were transformed into NEB Stbl cells and grown at 30C. Plasmid extraction from liquid cultures were performed using either Miniprep (Qiagen, #27106) or Midiprep (Macherey-Nagel, # 740412.50). All constructs were sequence verified either by sanger sequencing (Azenta) or whole plasmid sequencing (Quintara Biosciences and Plasmidsaurus).

### Prime Editing in HEK293T Cells

Prime editing experiments performed in HEK293T cells were performed by transfecting 16ug of pooled components encoded in plasmid DNA using lipofectamine 3000 (Invitrogen, #L300008) into 6-wells when cells were ∼70% confluent. Media was removed and replaced with fresh media 24 hours later.

### Lentiviral Preparation

Lentivirus containing various SCRIBBLE barcode designs (described below) was produced using a third-generation lentivirus packaging system: pMDLg/pRRE (Addgene, catalogue number 12251), pRSV-Rev (Addgene, 12253) and VSVG envelope (Addgene, 12259). Lenti-X 293T (TakaraBio, #632180) cells were grown in Dulbecco’s modified Eagle’s medium (DMEM) (Gibco, #11995-065) supplemented with 10% fetal bovine serum (FBS) (Gibco, #10437028) and 1% Pen Strep (Invitrogen, #15140-163). Upon reaching 70% confluence, we transfected the helped plasmids along with lentivral plasmid encoding our construct using Polyethylenimine (PEI) (Polysciences, #23966-1) at a 1:3 ratio of DNA:PEI. 24h after transfection, media was replaced with fresh culture media. 72h after transfection, media were collected, passed through a 0.45 µM filter (Corning, 431220), and then ultracentrifuged at 22,000g for 2 h. Pellets were resuspended in 100 µl of sterile phosphate-buffered saline (PBS) (Gibco, #10010031) overnight at 4 °C and then aliquoted and stored at −80 °C. Virus was titered by performing a dilution series on Lenti-X 293T cells and quantifying the fraction of GFP+ cells.

### Alginate Bacterial Culture System

We used an alginate culture system^43^ to maintain even abundance of plasmid library while growing in e. coli. We purified 500ng of ligation product from each cloning step using phenol:chloroform (ThermoScientific, #15593031) and resuspending in 1ul of DNAse/RNAse-Free Water (Invitrogen, #10977-015) and added this to 100ul of thawed DH10B cells on ice. We then incubated the bacteria/plasmid mix for 30 min on ice before gently transferring the mix to ice-chilled 1mm electroporation cuvettes (Fisher Scientific # BTX610). This was done by splitting the mix in half (53ul of the mix per cuvette). We then electroporated the cells (Biorad Gene Pulser Electroporator) using the following settings: 2.0kv, 200ohms, 25uF. Following electroporation, we quickly added 2ml of pre-warmed NEB 10-beta/Stable Outgrowth Media (NEB, # B9035SVIAL) and transferred bacteria to 37°C bacterial shaker where we cultured for 2hr shaking at 225rpm. We then created an alginate gel by mixing bacterial outgrown to a 25ml alginate (Sigma-Alrdich, # A1603-100MG)/Luria Broth (MilliporeSigma, # L3522) solution (final concentration 0.5%) containing 100ug/ml carbenicillin (ThermoFisherScientific, #10177012), 10mM CaCO_3_ (Sigma-Aldrich, #310034-100G) and 40ul of 37% HCL (MilliporeSigma, #320331). After combining components of this solution, the mixture is mixed by rocking for 15 min and then poured in a 150mm plate (CellTreat, #**229650**) which is then transferred to a 37C bacterial incubator. After 12-20hr of growth, the gel is melted by adding 6.25ml (1/4 volume) of a 20X SSC (ThermoFisherScientific, #15557044) and 40mM EDTA (ThermoFisherScientific, #AM9260G) solution and shaking at a room temperature for 15 min. Bacteria were then pelleted by centrifugation (5 min at 5000g), the supernatant was removed, and then the pellet rinsed another 6.25ml of melting solution. Bacteria were then resuspended in 150ml of LB with 100ug/ml carbenicillin containing 0.05mg/ml Alignate Lyase (MilliporeSigma, # A1603-100MG) and cultured in bacterial shaker for 4-8 hours to increase mass. Plasmids were then extracted using a standard midiprep kit (Macherey-Nagel, # 740412.50).

### Experimental Model and Subject Details

All animal experiments were conducted in compliance with the protocol IS00001679, approved by the Institutional Care and Use Committees (IACUC) at Harvard University. Experiments were performed on tissue collected from wild-type male and female CD1 mice (Charles River, Strain #22).

### In Vivo P0 Electroporation

*In vivo* electroporation was performed in postnatal day 0 (P0) CD1 mice (Charles River, Strain #22). Plasmids used in an experiment were combined and diluted to desired concentration using DNAse/RNAse-free water (Invitrogen, #10977-015) along with 10X PBS (Invitrogen, AM9625) diluted to 1X and 0.1% (w/v) of fast-green (Sigma-Alrich, F7252-5G). Needles were prepared by pulling glass capillaries (Drumond, # 5-000-2100) using a horizontal needle puller (Sutter Instrument Co., Model P-97; Heat: 645, Pull: 30, Vel: 45, Time:50) and then beveling the tip to 35° using a micropipette grinder (Narishige, #EG-401). Animals were anesthetized using ice and were confirmed to be sufficiently anesthetized by lack of toe pinch response in accordance with IACUC protocols. The eyelids of the electroporated eye were gently separated using a sharp 25g needle and a drop of Proparacaine Hydrochloride ophthalmic solution, USP 0.5% (Akorn, NDC 17478–263-12) was applied to the cornea. The eye was then gently lifted by retracting eyelids using Dumont AA forceps (Fine Scientific Tools, # 11210-10) and plasmid mix was delivered subretinally using a beveled glass needle attached to a Femto Jet Express Microinjector (Eppendorf, 5248000261, discontinued). The injected eye was then electroporated using the Super Electroporator (Nepagene, Nepa21 Type II) using the following settings: Voltage: 80V, Pulse Length: 50msec, Pulse Interval: 950msec, Pulses: 5, Decay Rate: 0, Polarity: +. Pups were placed on heated mat for recovery before being returned to their mother.

### Mouse Retinal Explant Electroporation

Intact globes were dissected from embryonic mice of the desired age using fine forceps (Fine Scientific Tools, #11252-30 and #11251-35) and transferred to a dissection dish (Scintica, DD-ECON-90) in room temperature PBS (Gibco, #10010031). Plasmid mix was then injected into subretinal space using glass needles prepared as described above using an aspirator tube assembly (MilliporeSigma, A5177). Immediately after injection, the whole globe was electroporated using the Super Electroporator (Nepagene, Nepa21 Type II) and tweezer electrodes (Bulldog Bio, #CUY650P2) using the following settings: Voltage: 40V, Pulse Length: 50msec, Pulse Interval: 950msec, Pulses: 5, Decay Rate: 0, Polarity: +.

### Mouse Explant Culture

Following retina electroporation, sclera and cornea were dissected away leaving intact retina, lens, and vitreous which were transferred to an autoclaved 1.7ml Microtube (Axygen, MCT-175-T) filled with 1ml of pre-warmed retina culture media made from DMEM (Gibco, #11995-065), F12 (Thermo Fisher Scientific, #11765054), 10% FBS (Gibco, #10437028), and 1U/ml Penicillin/Streptromycin (Invitrogen, #15140-163). Up to 4 retinas were pooled in a single tube.

Samples were placed on a variable speed rocker-shaker (VWR) in a 37°C, humidified warm-room for the duration of the experiment. Media was half-changed every day.

### In ovo Electroporation

Fertilized White Leghorn chicken eggs (Charles River) were transferred from cool room to warm room (38°) approximately 36-40 hours prior to the desired injection time so that embryos were at Hamburger Hamilton Stage 10/11. Prior to injection, 4-5ml of albumin was removed using a 18g needle to make a small hole in egg shell with care not to hit yolk. The top of the egg was removed using fine curved scissors (Fine Scientific Tools, #14061-11). Plasmid mix was injected into left optic vesicle using hand-pulled glass needles and aspirator tube assembly as previously described. Electroporation was performed using a Super Electroporator (Nepagene, Nepa21 Type II) and wire electrodes using the following settings: Voltage: 40V, Pulse Length: 50msec, Pulse Interval: 950msec, Pulses: 5, Decay Rate: 0, Polarity: +. The eggs were then re-sealed with Duck HD Clear Packing Tape (Amazon # B000GR5XCW) and returned to the warm room (38°C) until desired collection time.

### *In Silico* Identification of Putative Active Sites

To identify putative SCRIBBLE protospacer sequences (referred to as active sites in the manuscript) that were both predicted to support high rates of Cas9 editing and not found in genomes of a broad set of species, we performed a series of *in silico* screens. Full documentation of these screens is available upon request. In brief, we first identified all potential protospacer sequences not present in the genomes of the following species which were chosen because they were either a classic model organism and/or a species that might be of experimental interest: *Homo sapiens, Mus musculus, Rattus norvegicus, Macaca mulatta, Callithrix jacchus, Gorilla gorilla, Mustela putorius furo, Sus scrofa, Gallus gallus, Xenopus tropicalis, Danio rerio, Drosophila melanogaster, Caenorhabditis elegans*. Since the computational time of aligning all unique 20bp sequences (4^20 ≈ 1.1 trillion sequences) across multiple genomes while also allowing for mismatches is prohibitively long, we took an iterative approach where we aligned all combinations of shorter sequences, extending only those that didn’t align to the 13 genomes with progressively longer combinations of sequence until G + 19mer length sequences were tested (**Supplemental Figure 1E**). Previous studies have found that Cas9 complexes can pause on mismatched protospacers that have identical pam-proximal sequences^44^. We reasoned that although these short matches are often not sufficient to promote cleavage, they might serve as potential “sinks” where Cas9 complex is drawn towards and therefore unable to edit the desired protospacer sequence. Therefore, while building shorter sequences to align, we choose sequences with low abundance pam-proximal sequences (**Supplemental Figure 1F**). After performing this mapping strategy, we scored each unmapped sequences using CRISPOR^31^ to calculate the Doench Score^30^ (Doench 2016, Rule 2) for all novel sequences, using the endogenous HEK3 locus as flanking sequence and replacing the HEK3 protospacer sequence with each putative protospacer. At the time this analysis was performed, there were no prime editor calculators available, but we reasoned that this approach would identify sequences that were likely to be successfully edited by Cas9 which would be a surrogate for prime editing activity. We choose the top 998 scoring sequences along with the endogenous HEK3 and VEGFA sequences serving as controls to advance to the next screening stage.

### SCRIBBLE Screen 1 (Target)

To create a lentiviral screening library containing paired epegRNA and protospacer sequences, we designed an oligo pool containing Forward_Primer: Spacer_Sequence: Cloning_Site_1: RTT:Clonging_Site_2: Barcode:Clonging_Site_3: Protospacer:Reverse: Reverse Primer (Twist Biosciences). This pool was cloned downstream of a U6 promoter in a minimal plasmid. Next, a gRNA scaffold: Downstream “Flap” sequence was cloned into Cloning_Site_1. Then, an TevopreQ1 pseudoknot + PolyT (7) sequence was cloned into Cloning_Site_2. Then, 100bp of upstream flanking sequence from the endogenous HEK3 site was cloned into Cloning_Site_3, creating the final “shuttle” plasmid pool which contained a complete cassette of all paired epegRNA and target protospacer sequences necessary for this screen. The “acceptor” plasmid was created from a modified pSicoR lentiviral plasmid (Addgene, #11579) in which the U6/LoxP sequence was replaced with a cloning acceptor site followed by the endogenous downstream HEK3 flanking sequence. The complete cassette from the shuttle plasmid was then cloned into the shuttle vector forming the final lentiviral plasmid library. Throughout all stages of generating this library, we sought to maintain sequence diversity by performing large-scale transformation of cloned products into DH10B electrocompetent bacteria (NEB, #C3020K) and growing the transformants using an alginate culture system^43^ (see methods section). After generating the final plasmid library, we generated high-titer lentivirus (see methods section).

To perform the screen, we generated a pool of stably transduced HEK293T with ∼3500X coverage of each sequence by performing sparse viral labeling (MOI of ∼0.11) and used FACS is enrich for labeled cells. After allowing for several passages to recover from FACS, we plated transduced HEK293T cells in 10cm plates which served as technical replicates. When cultures were 70% confluent the following day, we transfected cells with a mix of plasmids using Lipofectamine 3000 (ThermoFisher Scientific, #L3000008) and Opti-Mem (ThermoFisher Scientific, #51985034) using the following mix:

- 48.6ug of CAG:PEmax (cloned from Addgene, #174820)
- 3.2ug of CAG:mCherry
- Optimem: 500ul
- Lipofectamine 3000: 43.4ul
- P3000: 29ul

Three days after transfection, we isolated mCherry+ cells using FACS and captured 3.16M cells in replicate 1 and 3.56M cells in replicate 2. After FACS, we centrifuged cells at 2000g for 10 min, removed supernatant and extracted DNA using the Quick-DNA Midiprep Plus Kit (Zymo, #D4075), eluting each sample in 47.8ul of elution buffer. We used 1ul of each sample to measure DNA concentration and split the remaining volume evenly across four separate reactions where we amplified lentiviral sequences using a custom strand-tagging approach. This approach labels each recovered molecule with a unique molecular identifier prior to amplification so that differences in PCR amplification don’t confound the results.^45^ This library generation protocol is performed using the following steps:

1. Strand Tag Round 1
2. Exonuclease I Digestion
3. Strand Tag Round 2
4. Exonuclease I Digestion
5. Library Amplification Part 1
6. Library Amplification Part 2

#### Strand Tag Round 1

Mix the following reagents in PCR tubes (USA Scientific, #1402-4700):

- 25ul NEB Q5 HS MasterMix (NEB, #M0494L)
- 0.8ul 50uM Strand-tagging primer 1 (Oligo6, **Table 1**)
- 11.7ul Genomic DNA
- 12.5ul Water

Place tubes on thermocycler (BioRad, C1000 Touch Thermocycler) and run program:

1. 72C, 3 min 2. 98C, 30 sec
2. 62C, 5 min
3. 72C, 1 min
4. 4C, hold

#### Exonuclease I Digestion

To remove un-used strand-tagging primer, add 1ul of Thermolabile Exonuclease I to reaction mix (NEB, #M0568L), mixed by pipetting and transfer back to thermocycler and run the program:

1. 37C, 5 min
2. 80C 1 min
3. 4C, hold

#### Strand Tag Round 2

Add the following reagents in order to reaction mix:

- 2.5ul NEB Q5 HS MasterMix (NEB, #M0494L)
- 1.0ul 50uM Strand-tagging primer 2 (Oligo7, **Table 1**)
- 0.5ul Water

Place tubes on thermocycler (BioRad, C1000 Touch Thermocycler) and run program:

1. 72C, 3 min
2. 98C, 30 sec
3. 62C, 5 min
4. 72C, 1 min
5. 4C, hold

#### Exonuclease I Digestion

Repeat as previously described

#### Library Amplification Part 1

Add the following reagents to reaction in order:

- 10ul NEB Q5 HS Master Mix (NEB, #M0494L)
- 2.5ul 10uM SCRIBBLE Screen 1 Forward Primer (Oligo8, **Table 1**)
- 2.5ul 10uM SCRIBBLE Screen 1 Reverse Primer (Oligo9, **Table 1**)
- 4.0ul Water

Place tubes on thermocycler (BioRad, C1000 Touch Thermocycler) and run program:

1. 98C, 30 sec
2. 98C, 10 sec
3. 65C, 10 sec
4. 72C, 15 sec
5. Repeat 2-4 15x
6. 72C, 1 min
7. 4C, hold

Clean amplified libraries by performing dual-sided (0.8X-0.6X) SPRI clean-up (Beckman Coulter, #B23318):

1. Add 30 ul of SPRI to amplified libraries
2. Mix by pipetting gently 10 times with pipette set to 40ul
3. Incubate at room temperature for 5 minutes.
4. Move tubes to High setting of magnetic separator (ThermoFisher Scientific, #NC1469069) for 3 minutes until solution clears.
5. Remove supernatant, being careful not to disrupt beads.
6. Add 200ul of 70% EtOH, let incubate for 30 seconds
7. Repeat steps 5-6
8. Move tube to Low setting of magnetic separator
9. Let air dry for 1 min
10. Remove from magnet, add 10ul DNAse/RNAse-Free water, incubate for 5 min
11. Place tube on High setting of magnetic separator for 3 min
12. Carefully transfer supernatant to new PCR tube.

#### Library Amplification Part 2

Pool 2ul from each of the 4 sub-libraries in each replicate, and mix the following reagents in a PCR tube:

- 8ul pooled Sub-library PCR product
- DNAse/RNAse-Free water
- 25ul NEB Q5 HS Master Mix (NEB, #M0494L)
- 2.5 ul 10uM Read 1 Indexing Primer (Oligo 10/11, **Table 1**)
- 2.5 ul 10uM Read 2 Indexing Primer (Oligo 12/13, **Table 1**)

Place tubes on thermocycler (BioRad, C1000 Touch Thermocycler) and run program:

1. 98C, 30 sec
2. 98C, 10 sec
3. 65C, 10 sec
4. 72C, 15 sec
5. Repeat 2-4 13x
6. 72C, 1 min
7. 4C, hold

Clean amplified libraries by performing dual-sided (0.8X-0.6X) SPRI clean-up (Beckman Coulter, #B23318) as previously described in Library Amplification Part 1. Elute in 20ul of DNAse/RNAse-Free water.

### SCRIBBLE Screen 2 (Pairwise)

If order to conduct the pairwise SCRIBBLE screen, first we generated a pool of oligos containing the following sequence: Forward_Primer: Protospacer: gRNA_Scaffold: RTT(Downstream_Flap: New_Protospacer: 4bp_BarcodeFragment): Primer_Binding_Site: TevopreQ1_Fragment. We amplified this pool using Oligos 14 and 15 (**Table 1**). We cloned the resulting pool into a modified pU6-pegRNA-GG-acceptor (Addgene, 132777) containing the full lengthTeveProQ1 and PolyT (7) sequence adjacent to the second BsaI site. We then electroporated this cloned plasmid library into DH10B cells as previously described and harvested plasmid library. To confirm cloning of all elements and calculate relative abundance of each epegRNA in the library, we generated a next-generation sequencing library using 10ng of plasmid library as template in a two-step strand-tagging approach similar to what is described in the SCRIBBLE Screen 1 methods section. Strand-tagging was performed as previously described using Oligo 14 (Strand-tag step 1) and Oligo15 (Strand-tag step 2). After both strand-tagging steps, we performed a two-stage amplification exactly as described before. We sequenced libraries using the Novaseq 6000 (Illumina) in a paired-end 150 format.

Nest, we generated a set of SCRIBBLE barcode sites containing each of the 12 putative protospacer sequences. These 12 sequences were sub-amplified out of the final lentiviral plasmid library used in Screen 1 using pairs of primers (Oligos 19-52, Table 1) in which the forward primer bound to the protospacer sequence and the reverse primer bound the downstream portion of the HEK3 flanking sequence resulting in a ∼200bp amplicon. These sequences were then individually ligated into the “acceptor” plasmid used in Screen 1 in the same site the shuttle plasmid in Screen 1 was added. We then made an equimolar pool of these 12 constructs and generated lentivirus (see methods for Lentivirus generation) which we then used to transduce a 10cm plate of HEK293T cells at high MOI (∼2.3). Three days after transduction, we collected ∼6M GFP+ cells by FACS and replated them in a 10cm cell culture dish to recover. Three days later, we plated 2.4M cells in two 6 cm plates which would serve as the technical replicates. The following morning when cultures were 70% confluent, we transfected cultures with the following plasmid mix using Lipofectamine 3000:

- epegRNA Pool, 5.5ug
- CAG:PEMax, 16.5ug
- 250ul Opti-Mem
- 16.5ul Lipofectamine 3000
- 22ul P3000

The following morning, culture media was replaced with fresh media. Two days later, we split the cultures 1:3 to prepare them for transfection the following day. We repeated this 4-day cycle for a total of 5 rounds before extracting DNA using the Quick-DNA Midiprep Plus Kit (Zymo, #D4075). Over the course of the entire experiment, we only kept 1 of the newly split plates meaning that we maintained at least 2.4M cells minimum per replicate throughout the screen. We then split DNA from each replicate into 16 equal sub-libraries and generated a strand-tagged NGS library using the following steps:

1. Strand-tag with Oligo 18 (**Table 1**) as previously described in Screen 1 methods
2. Exonuclease I Digestion as previously described in Screen 1 methods
3. PCR Amplification of each sublibrary with Oligos 19 and 20 using NEB Q5 HS MM as previously described in Screen 1 methods using the following PCR program:
  a. 98C, 30 sec
  b. 98C, 10 sec
  c. 65C, 10 sec
  d. 72C, 10 sec
  e. Rep b-d, 24 times
  f. 4C, hold
4. Strand-tag with in-line indexing primers (Oligos 21-28, **Table 1**) using one primer for two of the 16 different sub-libraries in each sample.
5. Exonuclease I Digestion as previously described in Screen 1 methods.
6. Pool 13ul of each library together forming 2 sets of 8 sublibraries for each technical replicate (with one sub-library strand-tagged with each of the 8 Oligos in step 4).
7. Clean libraries by performing dual-sided SPRI clean-up (1.2X-0.6X), elute in 20ul of DNAse/RNAse-free water.
8. Performing final PCR with IDT xGen UDI Primers Plate 1, (IDT, # 10005922) using PCR program:
  a. 98C, 30 sec
  b. 98C, 10 sec
  c. 65C, 10 sec
  d. 72C, 10 sec
  e. Rep b-d, 8 times
  f. 4C, hold
9. Perform 0.8X left-sided SPRI clean-up.

Libraries were sequenced on a Novaseq 6000 (Illumina) with paired-end sequencing 25 cycles on Read1 and 291 cycles on Read2.

### SCRIBBLE Edit Score

An edit score for each potential pair of sequential edits was calculated using the following equation, where N_Edit_ is the number of molecules carrying that edit, F_epegRNA_ is the frequency in the library of the epegRNA capable of making that particular edit, and N_Target_ is the number of molecules carrying the starting sequence.

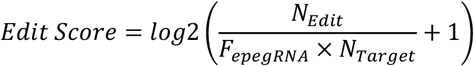

Adding 1 before the log transformation keeps the score finite for pairs at which no edits were observed. Only pairs in which the two protospacers differ were scored.

### SCRIBBLE Target Library Creation

To generate the SCRIBBLE Target plasmid library, we first generated an “entry” vector derived from the ampicillin resistance and origin of replication elements from the pU6-pegRNA-GG-acceptor plasmid^29^ (Addgene, #Plasmid #132777). To this, we added: piggyBac left (5’) inverted repeat, T7 RNA polymerase promoter, human U6 promoter, Twister TTRz, Broccoli aptamer, T3 RNA polymerase promoter, Barcode cloning site, SCRIBBLE downstream sequence, prime editor nicking site 3, 10x Genomics capture sequence 2, 100bp polyA, Twister TRz, U6 termination sequence, T7 terminator sequence, piggyBac right (3’) inverted repeat. While most of these elements were synthesized by Genescript, IDT, or Ansa, the Twister ribozyme system and broccoli sequences were cloned from a plasmid from Litke and colleagues^46^ (Addgene #124362).

We then generated a partially double-stranded sequence encoding the upstream primer site, a static barcode (VHDBx5) and either of the full-length SCRIBBLE protospacer sequences by annealing the following Oligos (**Table 1**, IDT):

- 2ul 100uM Oligo 1 or 2 (Active Site 2 or 7)
- 2ul 100uM Oligo 3 (Reverse Complement Upstream Primer)
- 2ul 100uM Oligo 4 or 5 (Reverse Complement Active Site 2 or 7)
- 5ul 10X T4 DNA ligase Buffer (NEB, #M0202T)
- 39ul H20

Annealing was performed on a thermocycler using the following program:

1. 98C, 5 min
2. -0.1C/cycle, 9 seconds
3. Repeat step 2 700x
4. 4C, hold

The insert was purified using the Zymo Clean and Concentrator kit (Zymo, #D4004) and eluted in 20ul of Elution Buffer. The SCRIBBLE Entry vector was linearized at the barcode cloning site using BsaI (NEB, # R3733S). The two components were then ligated using NEB T4 DNA Ligase (NEB, #M0202T) using 500ng of linearized backbone and 3x molar excess of insert in the following reaction:

- 500ng linearized SCRIBBLE Entry vector
- 3x molar excess annealed Barcoded SCRIBBLE insert
- 2ul 10X T4 DNA ligase (NEB, #M0202T)
- 1ul T4 DNA ligase (NEB, #M0202T)

The ligation was incubated at room temperature for 10 min and then for 16hours at 16C on a thermocycler. The following morning, the ligation reaction was cleaned using the Zymo Clean and Concentrator kit (Zymo, #D4004) and then eluted in 20ul of elution buffer. We then performed a negative selection digest to linearize undesired auto-annealed products by performing a 4hr digestion using BsaI (NEB, # R3733S). The reaction product was again cleaned with Zymo Clean and Concentrator kit and eluted in 6ul in elution buffer.

We then performed a large-scale electroporation of the barcoded SCRIBBLE target library using MegaX Electrocompetent e coli (ThermoFisher # C640003). In brief, we thawed 1 tube of MegaX cells on ice and then added all 6ul of eluted barcoded SCRIBBLE target library to the e. coli, keeping the tube on ice. We then incubated the bacteria/plasmid mix for 30 min on ice before gently transferring the mix to ice-chilled 1mm electroporation cuvettes (Fisher Scientific # BTX610). This was done by splitting the mix in half (53ul of the mix per cuvette). We then electroporated the cells (Biorad Gene Pulser Electroporator) using the following settings: 2.0kv, 200ohms, 25uF. Pre-warmed recovery media (ThermoFisher # C640003) was quickly added to cuvettes following electroporation and then transferred along with e coli to 14ml round bottom polystyrene tube (Falcon #352001) and incubated in a bacterial shaker at 37C, 225RPM for 1 hour. We then plated transformed bacteria across 4 large format Qtrays (Fisher NC9372402) with LB agar (Invitrogen 22700-041) containing 100ug/ml carbenicillin (Invitrogen #10177012) and incubated for 14 hours at 37C. We then added 20ml of LB to surface of each plate and scraped bacteria off the surface of plates using sterile cell scraper (Cell Treat #229305). Bacteria + LB was collected on ice, mixed by gentle pipetting and then processed in 6 midi preps per library (Macherey-Nagel NucleoBond Xtra Midi plus #740412.50). DNA was resuspended at a concentration of 10ug/ul.

### SCRIBBLE Barcode Recovery (From Bulk)

DNA was extracted using Quick-DNA Microprep Plus Kit (Zymo, #D4074) using manufacturer’s protocol. Strand-tagging was performed using 2X Q5 HS MM as described previously in this methods section, using a pool of Oligos 60-63. Samples were then treated with Thermolabile Exonuclease I (NEB, #M0568L) for 30 min at 37C and then at 90C for 1 min. Libraries were then directly amplified using Oligos 64-69 for 25 cycles and then cleaned using a 0.8X-left sided SPRI selection. Libraries were then amplified 8 more cycles by PCR using IDT xGen UDI Primers Plate 1, (IDT, # 10005922). Final libraries were cleaned with a 0.8X-left sided SPRI selection.

### SCRIBBLE Barcode cDNA Generation (From Bulk)

RNA was extracted from samples using Direct-zol RNA MiniPrep with TRI-Reagent (Zymo Research, #R2051) using manufacturer’s protocol. Reverse transcription was performed using Induro Reverse Transcriptase kit (NEB, #M0681L) using manufacturer’s protocol and a pool of Oligos 60-63 (**Table 1**). Samples were then treated with Thermolabile Exonuclease I (NEB, #M0568L) and RNAse A (Thermo Scientific, #EN0531) for 30 min at 37C and then at 90C for 1 min. Samples were then cleaned with 0.8X left-sided SPRI selection. Samples were then amplified by PCR with 2X Q5 HS MM (NEB, #M0494L) using manufacturer’s protocols and a pool of Oligos 64-68 and Oligo 69. Libraries were cleaned using a 0.8X-left sided SPRI selection and then amplified 8 more cycles by PCR using IDT xGen UDI Primers Plate 1, (IDT, # 10005922). Final libraries were cleaned with a 0.8X-left sided SPRI selection.

### scRNA-seq w/ SCRIBBLE

To prepare SCRIBBLE-labeled retinas for scRNA-seq, we dissociated three retinas by incubating in papain (Worthington, #LK003150) for 30 minutes at 37C. We then gently triturated samples with a P1000 pipeteman 20 times and then spun in centrifuge for 5 min at 300g. We then carefully removed supernatant, being sure to not disturb cell pellet. We then re-suspended cells in ovo-mucoid/DNAse I mixture as kit indicated, gently triturating 20 times using a P200 pipeteman. We again centrifuged cells for 5 min at 300g. We then resuspended cells in modified cell culture media diluted to 3% FBS using 1:1 mix of DMEM and F12. We then enriched for GFP+ cells using a On-Chip Sort microfluidics device. Sorted cells were then concentrated by centrifugation for 10 min at 500g. Cells were then captured using GEM-X Universal 3’ Gene Expression v4 (10X genomics, #1000686). During library preparation, we sub-amplified SCRIBBLE barcode reads using by modifying the CaRPool-seq^47^ protocol to target an upstream sequence in the SCRIBBLE barcode (Oligos 53-57, **Table 1**). As is detailed in the CaRPool-seq protocol, we retained the supernatant from the 0.6X SPRI selection performed after the PCR and purified it first with a 1.2X left-sided SPRI clean-up, and then a 1.0X left-sided SPRI clean-up, eluting in 20ul of Buffer EB. We then used a fraction of this smaller fragment cDNA library in an indexing PCR to make the libraries compatible with Illumina sequencing:

- 25ul 2X Q5 HS MM
- 2.5ul F Primer (Oligo 58)
- 2.5ul R Primer (Oligo 59)
- 2.0ul amplified cDNA
- 18ul water

PCR amplification was performed using the following program:

1. 98 C 30 sec
2. 98 C 10 sec
3. 65 C 20 sec
4. 72 C 20 sec
5. Repeat 2-4 7x (8-20 total cycles) depending on results
6. 4 Hold

The PCR was then cleaned using a 0.8X left-sided SPRI selection and eluted in 15ul of water. Libraries were sequenced on Illumina Novaseq 6000 sequencer using a 300 cycle kit with Read1: 28 cycles, Index1:10 cycles, Index2:10 cycles, Read2: 298 cycles. The transcriptome was aligned using 10X Cellranger and the three distinct specimens were identified based off single nucleotide polymorphisms using Cellsnp-lite^48^ and Vireo^49^.

## ACKNOWLEDGEMENTS

We thank members of the Cepko, Tabin, and Dymecki Labs for helpful discussions. We thank Ashley Hurst and the BPF Genomics Core Facility at Harvard Medical School (RRID:SCR_007175) for their expertise and instrument availability that supported this work.

## AUTHOR CONTRIBUTIONS

R.N.D. conceived the project, designed the method, conducted experiments, and led experimental analyses. Y.S. developed SCRIBBLE barcode bioinformatic pipeline. R.E.A conducted experiments including scRNA-seq and provided intellectual contributions. C.L. conducted experiments including single cell sorting and provided intellectual contributions. H.L.B. conducted experiments including explant electroporation and provided intellectual contributions. A.A.S. and D.R.L. provided insights to prime editing implementation. J.L.D., S.P., and D.R.L. produced the JLD1907 prime editor variant. E.W. performed in vivo P0 mouse electroporation. C.E.G. assisted in molecular cloning. C.L.C. supervised the project. R.N.D. and C.L.C. wrote the manuscript. All of the authors read and approved the final manuscript.

## ETHICS STATEMENT

All animal work was performed under protocols approved by Harvard Medical School.

## DATA AND CODE AVAILABILITY

Data and code will be available upon publication or by reasonable request.

## REFERENCES

1 Masland, R. H. The neuronal organization of the retina. Neuron 76, 266–280 (2012). 10.1016/j.neuron.2012.10.002

2 Shekhar, K. et al. Comprehensive Classification of Retinal Bipolar Neurons by Single-Cell Transcriptomics. Cell 166, 1308–1323 e1330 (2016). 10.1016/j.cell.2016.07.054

3 Li, J. et al. Comprehensive single-cell atlas of the mouse retina. iScience 27, 109916 (2024). 10.1016/j.isci.2024.109916

4 Turner, D. L., Snyder, E. Y. & Cepko, C. L. Lineage-independent determination of cell type in the embryonic mouse retina. Neuron 4, 833–845 (1990). 10.1016/0896-6273(90)90136-4

5 Williams, R. W. & Goldowitz, D. Structure of clonal and polyclonal cell arrays in chimeric mouse retina. Proc Natl Acad Sci US A 89, 1184–1188 (1992). 10.1073/pnas.89.4.1184

6 Hafler, B. P. et al. Transcription factor Olig2 defines subpopulations of retinal progenitor cells biased toward specific cell fates. Proc Natl Acad Sci US A 109, 7882–7887 (2012). 10.1073/pnas.1203138109

7 Suzuki, S. C. et al. Cone photoreceptor types in zebrafish are generated by symmetric terminal divisions of dedicated precursors. Proc Natl Acad Sci US A 110, 15109–15114 (2013). 10.1073/pnas.130355111O

8 Rompani, S. B. & Cepko, C. L. Retinal progenitor cells can produce restricted subsets of horizontal cells. Proc Natl Acad Sci US A 105, 192–197 (2008). 10.1073/pnas.0709979104

9 Bushnell, H. L. & Cepko, C. L. Distinct neurogenic progenitor cell populations balance cell type production in the embryonic mouse retina. bioRxiv (2026). 10.64898/2026.08.03.738451

10 Turner, D. L. & Cepko, C. L. A common progenitor for neurons and glia persists in rat retina late in development. Nature 328, 131–136 (1987). 10.1038/328131ao

11 Moody, S. A., Chow, I. & Huang, S. Intrinsic bias and lineage restriction in the phenotype determination of dopamine and neuropeptide Y amacrine cells. J Neurosci 20, 3244–3253 (2000). 10.1523/JNEUROSC.20-09-03244.2000

12 Huang, S. & Moody, S. A. Asymmetrical blastomere origin and spatial domains of dopamine and neuropeptide Y amacrine subtypes in Xenopus tadpole retina. J Comp Neurol 360, 442–453 (1995). 10.1002/cne.903600306

13 Gomes, F. L. et al. Reconstruction of rat retinal progenitor cell lineages in vitro reveals a surprising degree of stochasticity in cell fate decisions. Development 138, 227–235 (2011). 10.1242/dev.059683

14 He, J. et al. How variable clones build an invariant retina. Neuron 75, 786–798 (2012). 10.1016/j.neuron.2012.06.033

15 Kolb, H., Fernandez, E., Jones, B. W. & Nelson, R. About/FAQ. Webvision: The Organization of the Retina and Visual System (1995).

16 Yan, W. et al. Mouse Retinal Cell Atlas: Molecular Identification of over Sixty Amacrine Cell Types. J Neurosci 40, 5177–5195 (2020). 10.1523/JNEUROSC.0471-20.2020

17 Shekhar, K., Whitney, I. E., Butrus, S., Peng, Y. R. & Sanes, J. R. Diversification of multipotential postmitotic mouse retinal ganglion cell precursors into discrete types. Elife 11 (2022). 10.7554/elife.73809

18 Clark, B. S. et al. Single-Cell RNA-Seq Analysis of Retinal Development Identifies NFI Factors as Regulating Mitotic Exit and Late-Born Cell Specification. Neuron 102, 1111–1126 e1115 (2019). 10.1016/j.neuron.2019.04.01O

19 Lu, Y. et al. Single-Cell Analysis of Human Retina Identifies Evolutionarily Conserved and Species-Specific Mechanisms Controlling Development. Dev Cell 53, 473–491 e479 (2020). 10.1016/j.devcel.2020.04.009

20 Wu, F. et al. Single cell transcriptomics reveals lineage trajectory of retinal ganglion cells in wild-type and Atoh7-null retinas. Nat Commun 12, 1465 (2021). 10.1038/s41467-021-21704-4

21 Balasubramanian, R. et al. Phase transition specified by a binary code patterns the vertebrate eye cup. Sci Adv 7, eabj9846 (2021). 10.1126/sciadv.abj9846

22 West, E. R. et al. Spatiotemporal patterns of neuronal subtype genesis suggest hierarchical development of retinal diversity. Cell Rep 38, 110191 (2022). 10.1016/j.celrep.2021.110191

23 Holt, C. E., Bertsch, T. W., Ellis, H. M. & Harris, W. A. Cellular determination in the Xenopus retina is independent of lineage and birth date. Neuron 1, 15–26 (1988). 10.1016/0896-6273(88)90205-x

24 Xu, Q., Tam, M. & Anderson, S. A. Fate mapping Nkx2.1-lineage cells in the mouse telencephalon. J Comp Neurol 506, 16–29 (2008). 10.1002/cne.21529

25 Brzezinski, J. A. t., Kim, E. J., Johnson, J.E. & Reh, T. A. Ascl1 expression defines a subpopulation of lineage-restricted progenitors in the mammalian retina. Development 138, 3519–3531 (2011). 10.1242/dev.064006

26 McKenna, A. et al. Whole-organism lineage tracing by combinatorial and cumulative genome editing. Science 353, aaf7907 (2016). 10.1126/science.aaf7907

27 Chow, K. K. et al. Imaging cell lineage with a synthetic digital recording system. Science 372 (2021). 10.1126/science.abb3099

28 Chan, M. M. et al. Molecular recording of mammalian embryogenesis. Nature 570, 77–82 (2019). 10.1038/s41586-019-1184-5

29 Anzalone, A. V. et al. Search-and-replace genome editing without double-strand breaks or donor DNA. Nature 576, 149–157 (2019). 10.1038/s41586-019-1711-4

30 Doench, J. G. et al. Optimized sgRNA design to maximize activity and minimize off-target effects of CRISPR-Cas9. Nat Biotechnol 34, 184–191 (2016). 10.1038/nbt.3437

31 Concordet, J. P. & Haeussler, M. CRISPOR: intuitive guide selection for CRISPR/Cas9 genome editing experiments and screens. Nucleic Acids Res 46, W242–W245 (2018). 10.1093/nar/gky354

32 Chen, B. et al. Dynamic imaging of genomic loci in living human cells by an optimized CRISPR/Cas system. Cell 155, 1479–1491 (2013). 10.1016/j.cell.2013.12.001

33 Nelson, J. W. et al. Engineered pegRNAs improve prime editing efficiency. Nat Biotechnol 40, 402–410 (2022). 10.1038/s41587-021-01039-7

34 Doman, J. L. et al. Phage-assisted evolution and protein engineering yield compact, efficient prime editors. Cell 186, 3983–4002 e3926 (2023). 10.1016/j.cell.2023.07.039

35 Li, X. et al. Chromatin context-dependent regulation and epigenetic manipulation of prime editing. Cell 187, 2411–2427 e2425 (2024). 10.1016/j.cell.2024.03.020

36 Young, R. W. Cell differentiation in the retina of the mouse. Anat Rec 212, 199–205 (1985). 10.1002/ar.1092120215

37 Choi, J. et al. A time-resolved, multi-symbol molecular recorder via sequential genome editing. Nature 608, 98–107 (2022). 10.1038/s41586-022-04922-8

38 Loveless, T. B. et al. Open-ended molecular recording of sequential cellular events into DNA. Nat Chem Biol 21, 512–521 (2025). 10.1038/s41589-024-01764-5

39 Yu, Q. et al. In vivo reconstruction of the cell lineage history of a developing mouse with DNA Typewriter, from zygote to late organogenesis. bioRxiv (2026). 10.64898/2026.07.29.741625

40 Koblan, L. W. et al. High-resolution spatial mapping of cell state and lineage dynamics in vivo with PEtracer. Science 390, eadx3800 (2025). 10.1126/science.adx3800

41 Colgan, W. N. et al. Comprehensive Lineage Tracing Maps the Landscape of Cell Fate Decisions in Mouse Embryogenesis. bioRxiv (2026). 10.64898/2026.05.07.722278

42 Winter, E., Emiliani, F., Cook, A., Abderrahim, A. & McKenna, A. BASELINE: a CRISPR base editing platform for mammalian-scale single-cell lineage tracing. Nucleic Acids Res 54 (2026). 10.1093/nar/gkag769

43 Rabe, B. A. Investigation of the Multiple Roles of Notch Signaling in Embryonic Retinal Development Using Novel High-Throughput Techniques, Harvard University Graduate School of Arts and Sciences, (2020).

44 Wu, X. et al. Genome-wide binding of the CRISPR endonuclease Cas9 in mammalian cells. Nat Biotechnol 32, 670–676 (2014). 10.1038/nbt.2889

45 Xing, D., Tan, L., Chang, C.H., Li, H. & Xie, X. S. Accurate SNV detection in single cells by transposon-based whole-genome amplification of complementary strands. Proc NatlAcad Sci USA 118 (2021). 10.1073/pnas.2013106118

46 Litke, J. L. & Jaffrey, S. R. Highly efficient expression of circular RNA aptamers in cells using autocatalytic transcripts. Nat Biotechnol 37, 667–675 (2019). 10.1038/s41587-019-0090-6

47 Wessels, H. H. et al. Efficient combinatorial targeting of RNA transcripts in single cells with Cas13 RNA Perturb-seq. Nat Methods 20, 86–94 (2023). 10.1038/s41592-022-01705-x

48 Huang, X. & Huang, Y. Cellsnp-lite: an efficient tool for genotyping single cells. Bioinformatics 37, 4569–4571 (2021). 10.1093/bioinformatics/btab358

49 Huang, Y., McCarthy, D. J. & Stegle, O. Vireo: Bayesian demultiplexing of pooled single-cell RNA-seq data without genotype reference. Genome Biol 20, 273 (2019). 10.1186/s13059-019-1865-2

